# Unfold to refold: Tracking the initial steps of PrP^C^ unfolding in the context of PrP^Sc^ propagation

**DOI:** 10.1101/2025.10.11.681810

**Authors:** Sanaz Sabzehei, Marta Rigoli, Raúl Cacheiro, Iria Díaz-Arias, Hasier Eraña, Rubén P. Lago, Arcadio Guerra, Human Rezaei, Joaquín Castilla, Emiliano Biasini, Víctor M. Sánchez-Pedregal, Manuel Martín-Pastor, Jesús R. Requena

**Affiliations:** CIMUS Biomedical Research Institute and Department of Medical Sciences, University of Santiago de Compostela-IDIS, Spain; Department of Cellular, Computational and Integrative Biology, University of Trento, Povo, TN, Italy; CIC BioGUNE, Basque Research and Technology Alliance (BRTA), Prion Research Lab, Derio, Spain; ATLAS Molecular Pharma S. L. Bizkaia Technology Park, Derio, Spain; Centro de Investigación Biomédica en Red de Enfermedades Infecciosas (CIBERINFEC), Carlos III National Health Institute, Madrid, Spain; CIQUS Research Institute, University of Santiago de Compostela, Spain; Université Paris-Saclay, INRAE, UVSQ, VIM, Jouy-en-Josas, France; IKERBASQUE, Basque Foundation for Science, Bilbao, Spain; Department of Organic Chemistry, University of Santiago de Compostela, Spain; Unidade de Resonancia Magnética, CACTUS, University of Santiago de Compostela, Spain

**Keywords:** Prion, PrP^Sc^, PrP^Sc^ propagation, PrP^C^, PrP^C^ unfolding, PrP^C^ NMR, MD

## Abstract

It might have been believed that elucidation of the atomistic structure of PrP^Sc^ would lead to an immediate understanding of the mechanism of prion propagation. However, PrP^Sc^, now known to be a “simple” amyloid, can just template a previously unfolded polypeptide chain. Therefore, PrP^Sc^ can easily template the disordered ∼90-120 domain of an incoming PrP^C^ molecule, but not its ∼121-231 folded domain (FD). The FD needs to accommodate into the ∼121-230 PrP^Sc^ surface, an inert “procrustean bed”. Thus, a mechanism for concerted unfolding/refolding of the FD must exist, with FD unfolding as a key element. To explore how this might happen, we performed thermal unfolding of recombinant bank vole PrP^C^(90-231), that is a universal PrP^Sc^ propagator PrP^Sc^, tracking changes at the residue level with solution NMR to pinpoint early unfolding propensity. Our data suggest that a key early event is the destabilization of the short β1-β2 assembly, and that the segment contiguous to the disordered tail, ∼121-140, encompassing β1 and its adjacent coils, is the most likely region to unfold first. Spectroscopic data obtained at higher temperatures suggest that portions of alpha helix α2 are likely the last elements of the FD to unfold and refold into the PrP^Sc^ conformation. Molecular Dynamics simulations assisted the interpretation of these changes and suggest separation of α1 from the rest of the FD ensemble. Our data provide a conceivable timeline of the early events in PrP^Sc^-assisted conversion of PrP^C^ and should serve as a starting framework to develop a future atomistic model of PrP^Sc^ propagation.

## Main

Nearly three decades after the atomic structure of the cellular prion protein (PrP^C^) was elucidated (1), the structure of its pathogenic counterpart, its scrapie isoform PrP^Sc^, has finally been resolved (2). This breakthrough, subsequently confirmed across multiple prion strains (3–5), marks a pivotal moment in prion research. It opens the door to understanding the molecular basis of prion propagation, defined as the conversion of PrP^C^ into PrP^Sc^, facilitated by pre-existing PrP^Sc^ molecules.

Propagation is one of the two defining features of a prion (6). The other is its ability to transmit between hosts, typically from the brain of an affected individual to that of a new host. In natural settings, and in the absence of human intervention, inter-host spread usually occurs via the oral route, and its mechanisms have been well elucidated (7–9). Therefore, the main challenge remaining to fully understand the PrP^Sc^ prion is determining how it catalyzes the conformational conversion of PrP^C^ into the alternatively folded PrP^Sc^ conformation, *i.e.*, into a likeness of itself.

Contemplation of PrP^C^ and PrP^Sc^ immediately suggests the first step of conversion: the N-terminal ∼23-120 domain of PrP^C^ is intrinsically disordered and the corresponding ∼90-120 domain of the templating surface of PrP^Sc^ (2–5) offers an array of unpaired - NH and -CO moieties that are highly predisposed to form hydrogen bonds with -CO and -NH groups in equivalent residues of an incoming PrP^C^ molecule. The decrease in entropy resulting from the transition from a disordered to a cross-β conformation is generously offset by the higher decrease in enthalpy resulting from the extensive array of hydrogen bonds in the newly molded Parallel-In-Register-Beta-Sheet (PIRIBS) segment (2). This well-known mechanism, commonly referred to as “seeded amyloid propagation”, is driven by specific molecular interactions often described as “lock and dock” events (10, 11).

Once the disordered ∼90-120 segment of PrP^C^ has adopted the PrP^Sc^ conformation, the remaining FD (residues ∼121-230) presents a more complex challenge. This globular region of PrP^C^ comprises three α-helices, two short antiparallel β-strands, and several well-defined connecting loops (1, 7, 12). The corresponding region in PrP^Sc^ is structurally inert: it lacks dynamic features, and must function primarily as a passive template. Its role would be limited to capturing segments of PrP^C^ that have already undergone partial unfolding. Such templating surface can be likened to a “procrustean bed” (13), requiring the incoming FD to conform precisely to its extended topology. To achieve this, the FD must first undergo unfolding as a prerequisite for its subsequent refolding into the PrP^Sc^ structure (see Suppl. Fig. S1). We envision that initially the FD likely remains suspended facing the templating surface, unable to engage until structural rearrangement occurs.

Importantly, the FD is a compact, roughly spherical domain composed of two subdomains: β1-α1-β2 and α2-α3, connected by a flexible hinge region spanning residues ∼164-168 (1, 12; Suppl. Fig. S2). For successful templating, these subdomains must separate and align with the extended N- and C-terminal lobes of the PrP^Sc^ surface (2, 14).

A substantial body of research has investigated the unfolding behavior of PrP^C^ (15–25). Most of these studies have employed elevated temperatures to accelerate unfolding, a process that naturally occurs over days, so that it can be observed within minutes. However, these investigations primarily focus on exploring and modelling spontaneous amyloid formation, which differs from PrP^Sc^-templated conversion even though both processes likely share some mechanistic elements. It is noteworthy that amyloids generated *in vitro* under conditions of simple PrP *misfolding*, *v.g*., shaking under denaturing conditions, have structures with short C-terminal PIRIBS cores that are much smaller than those of *bona fide* PrP^Sc^ (26, 27). We are specifically interested in the very early events of *unfolding*, before partially unfolded PrP^C^ conformers enter a pathway of spontaneous generation of PrP amyloids that are different from PrP^Sc^. In other words, we are interested in identifying early events within the PrP^C^ FD that result in partially unfolded conformations that can be trapped by the templating PrP^Sc^ surface in the context of PrP^Sc^ propagation.

Here we describe studies aimed at identifying such early unfolding events, using solution NMR and MD simulations. We compare our data with results from earlier studies and build a tentative timeline of early FD unfolding, which should be a first step for our ultimate aim: building an atomistic model of PrP^Sc^ propagation.

## Results

### Thermal unfolding of BVPrP^C^(90–231) is reversible up to 45 °C and then becomes irreversible

We analyzed the thermal unfolding of recBVPrP^C^(90–231) by monitoring the reduction in the α-helix-associated 222 nm minimum in the far-UV CD spectrum as temperature increased. As shown in Fig. 1A, the unfolding followed a two-state Boltzmann sigmoidal model (Eq. 8), with no detectable intermediate and a melting temperature (Tm) of 63 °C.

**Fig. 1.**
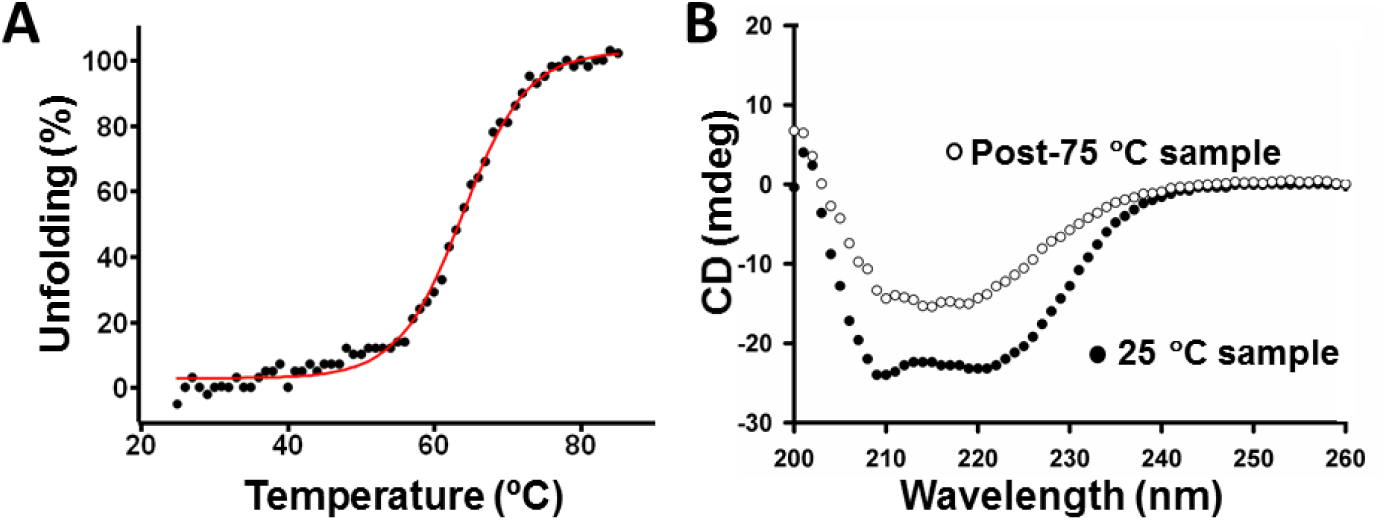
CD analysis of temperature-induced unfolding of BVPrP^C^(90–231). *(A)* Unfolding curve evaluated as decrease in CD signal at 222 nm (α-helix content) over temperature; the datapoints were fitted to the Boltzmann sigmoidal function (Eq. 8, continuous red line). The fitted parameters were A1=2.78, A2=102.67, x0=63.80, dx=4.41. *(B)* Comparison of CD spectra at the beginning (25°C, fresh sample) and after heating at 75°C and cooling down to 25°C.

To assess reversibility, a sample was heated to 75 °C for 1 hour and then cooled to 25 °C. The resulting CD spectrum (Fig. 1B) showed a ∼50% decrease in overall intensity and a shift toward a β-sheet-rich profile. Notably, the sample remained fully soluble throughout, with no precipitation observed even after centrifugation at 16,000g for 15 minutes.

We then employed solution NMR spectroscopy to obtain detailed, residue-specific insights into the thermal unfolding fingerprint. First, a heteronuclear single quantum coherence (HN-HSQC) spectrum of a sample of freshly folded ^13^C,^15^N-labeled recBVPrP^C^(90–231) dissolved in 10 mM sodium acetate buffer at pH 5, was recorded at 25 °C. Peak assignment was performed by comparison with data published by Christen *et al*. for BVPrP^C^(121–231) (12) and deposited in the BioMagResBank (www.bmrb.wisc.edu), accession numbers 15824, 15845). Minor spectral differences were observed due to the pH difference between our sample (pH 5) and Christen’s (pH 4.5), as well as the presence of the 90-120 amino acid segment in our construct. To resolve ambiguities arising from these differences and overlapping peaks, we conducted triple resonance experiments, enabling unambiguous assignment of uncertain signals (Suppl. Fig. S3). Resonances corresponding to the folded domain (residues 121-231) exhibited chemical shifts characteristic of structured proteins, while those from the 90-120 region displayed chemical shifts typical of disordered domains (Suppl. Fig. S3).

We subsequently recorded a series of N-HSQC spectra at temperatures ranging from 15 °C to 75 °C to investigate temperature-dependent structural changes in recBVPrP^C^(90–231). Up to 45 °C, the spectra retained strong signal intensities, narrow peak widths, and well-dispersed chemical shifts. Peak positions shifted gradually (Fig. 2 and Suppl. Fig. S4), which is consistent with the known temperature dependence of protein amide proton chemical shifts (28).

**Fig. 2.**
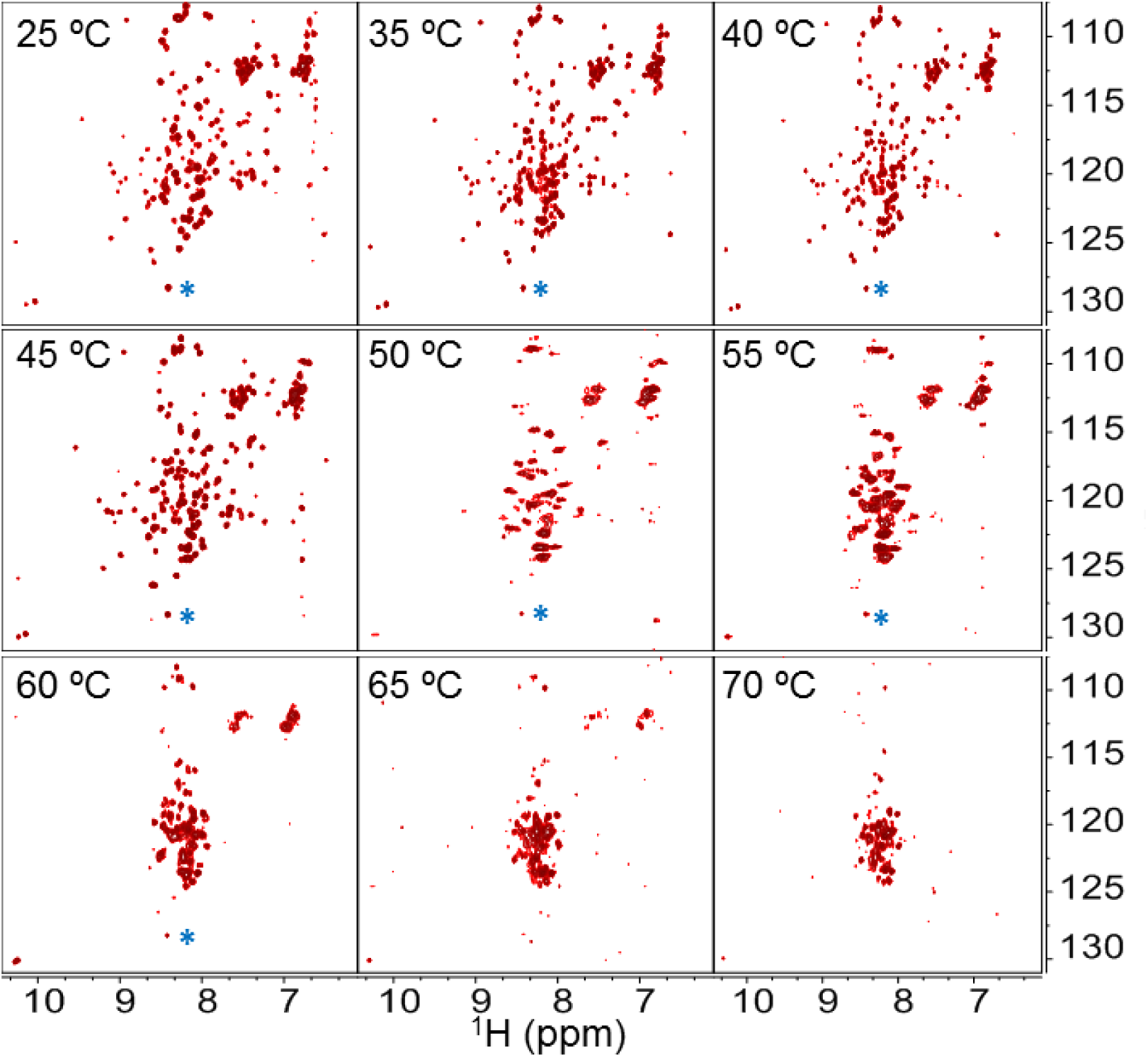
Temperature-induced unfolding of BVPrP^C^(90–231) tracked with solution NMR. N-HSQC spectra of the backbone amides from 25 to 70 °C. Each spectrum was recorded at the indicated temperature. The chemical shifts were referenced to the signals marked with the asterisk to facilitate comparison. The spectra are represented at the same noise level.

However, beginning at ∼50 °C, notable spectral changes emerged. Peaks began to broaden, and several signals diminished in intensity or disappeared entirely (Fig. 2). These alterations became more pronounced with further temperature increases.

To evaluate the reversibility of these changes, each sample was cooled back to 25 °C after heating. One sample per temperature condition was analyzed. Remarkably, samples heated to ≤ 45 °C recovered their original 25 °C N-HSQC spectra upon cooling, indicating that structural changes up to this temperature are reversible. In contrast, samples heated above 50 °C exhibited persistent spectral alterations after cooling (Suppl. Fig. S5), suggesting that structural changes occurring beyond this threshold are irreversible. Moreover, the broadening or even disappearance of the peaks suggests a polymerization/aggregation process which is consistent with the irreversible structural change observed by the CD thermal unfolding experiments performed at lower protein concentration (Fig. 1).

### Temperature dependence of amide H-N chemical shifts suggests early BVPrP^C^(90–231) unfolding events

Given that BVPrP^C^(90–231) remains structurally reversible up to 45 °C under our experimental conditions, we examined residue-level dynamic changes within the 15-45 °C temperature range. To achieve this, we used three classic solution NMR methods targeting backbone amide NH signals. Each of these methods offers complementary information regarding the “mobility” (used here in a broad sense) of individual residues. Together, they support a tentative mapping of regions within the FD that may be predisposed to initiate partial unfolding (Suppl. Fig. S6).

More precisely, we analysed: (a) ^15^N relaxation parameters (heteronuclear NOE and R2/R1 relaxation rates); (b) amide temperature coefficients Tc; and (c) the non-linearity of Combined Chemical Shift Differences (CCSD) of amide HN groups.

a. ^15^N relaxation parameters: We quantified longitudinal (R_1_), transverse (R_2_) relaxation rates, and steady-state ^15^N{^1^H} NOE values. The R_2_/R_1_ ratio and ^15^N NOE are sensitive reporters of the effective correlation time in the picosecond-to-nanosecond timescale experienced by each backbone NH group (29). Residues exhibiting low R_2_/R_1_ ratios or low ^15^N NOE values are interpreted as highly flexible or dynamically disordered, whereas higher values suggest rigidity with a higher contribution of slower motions or slow conformational exchange (30, 31).
b. Amide HN Temperature Coefficients (Tc): From the temperature dependence of HN proton chemical shifts we derived their Tc values, which provide insight into the stability of hydrogen bonds and solvent exposure of the HN. Less stable hydrogen bonding typically associates with backbone mobility.
c. CCSD analysis of the amide proton HN and its directly attached ^15^N (16): We examined the temperature-dependent changes of the chemical shifts of NH peaks in N-HSQC spectra. Analysis of the deviations from linearity of the CCSD identifies residues that explore alternative conformations in the fast-exchange regime.

It is important to note that, within the temperature range used for Tc and CCSD studies in this work, all backbone amide signals were in fast conformational exchange in the chemical shift timescale.

As seen in Fig. 3A,B, two clear segments can be distinguished in BVPrP^C^(90–231) according to their ^15^N NOE outputs at the two temperatures tested (25 and 45 °C): the ∼90-120 segment displays negative values with the exception of a stretch of residues with positive values at its middle (∼Q98-N108). In contrast, the ∼121-231 segment is characterized by positive values with a few exceptions. These results agree with previous data reported for and human PrP^C^(90–230), mule deer PrP^C^(90–231) or Syrian hamster PrP^C^(90–230) (15, 32, 33). Also in agreement with these studies, the border between these two disordered/ordered subdomains is characterized by a group of residues (∼V121-G126) with values that cross the zero line and slowly and steadily increase until reaching positive values of the same magnitude as the mean value of FD residues. These residues are therefore neither fully ordered nor disordered, showing increasing rigidity from the N to C ends of the ∼V121-G126 segment.

**Fig. 3.**
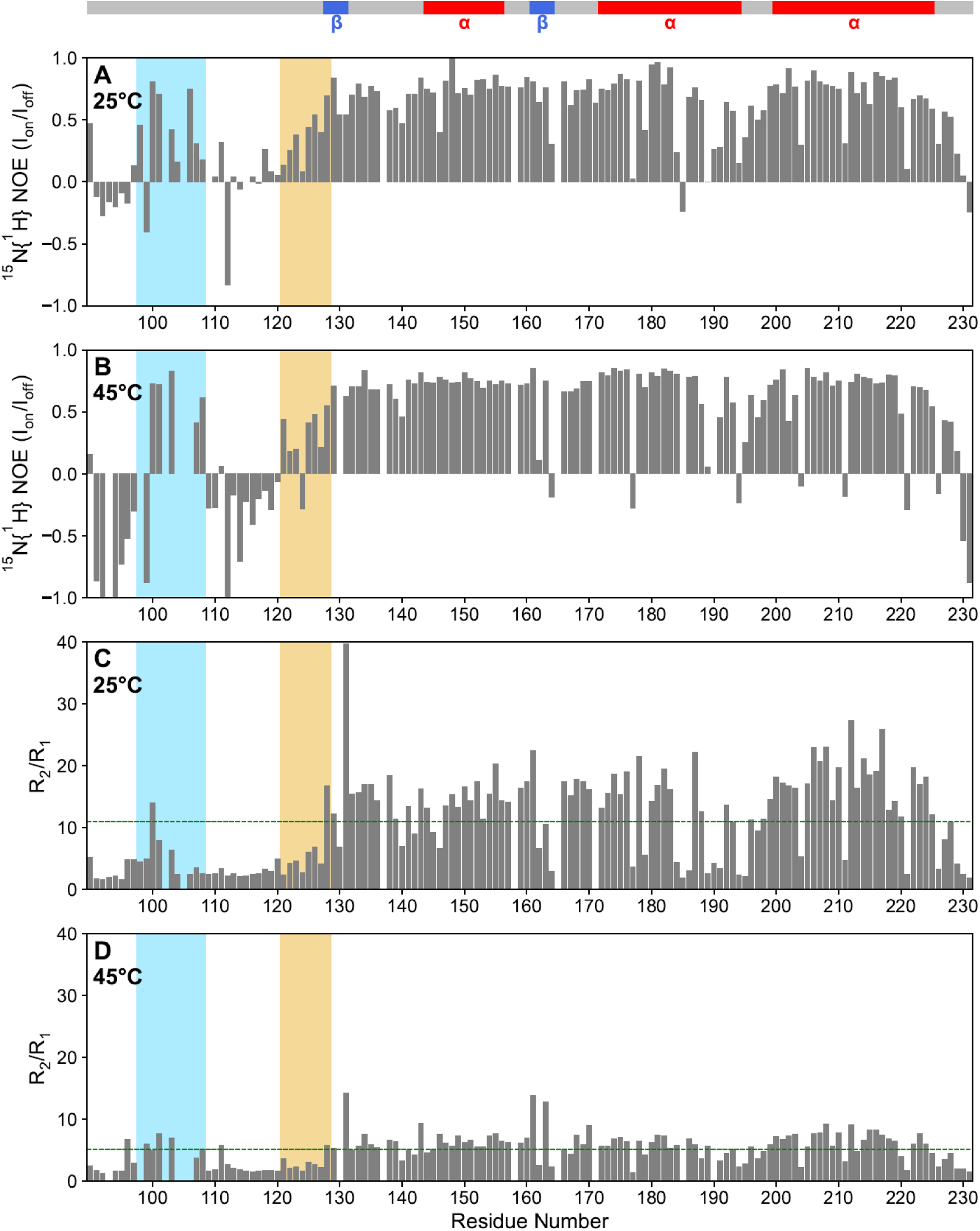
^15^N-NMR relaxation analysis of BVPrP^C^(90–231). ^15^N NOE at *(A)* 25 °C and *(B)* 45 °C and R2/R1 ratios at *(C)* 25 °C and (*D)* 45 °C. A transition zone with increasing ^15^N NOEs and R2/R1 values, respectively, corresponding to the ∼121-128 stretch is conspicuous (orange rectangles). This sequence is located at the border between the disordered 90-120 and fully folded ∼129-231 domains (see main text). Blue rectangles mark a stretch of residues within the disordered tail that exhibit unusual ^15^N NOEs and R2/R1 values, suggestive of transient local order.

R_2_/R_1_ plots show also conspicuous signatures of the two subdomains according to their group dynamics, faster and slower mobilities. Residues in 90-120 exhibit very low R_2_/R_1_ values (<5), indicative of fast internal motions, whereas residues within the ∼128-231 segment have values in the 10-20 range, typical of well folded domains. As seen for ^15^N NOE, a transition zone with increasing values is seen for the residues at the border between these two segments (∼121-126) (Fig. 3C,D). Residue G131, located at the end of β1, exhibits a remarkably high positive R_2_/R_1_ at 25 °C that paradoxically could indicate fast motions and/or conformational exchange (*i.e.* extreme flexibility). At 45 °C the same residue G131 together with V161 and Y163 located in β2 also exhibit the abnormally highest R_2_/R_1_ ratios.

We next calculated the Tc values of those backbone amide proton signals that showed linear behavior in the N-HSQC spectra over the 15-45 °C temperature range (*vide infra*). The amide Tc value largely reflects the stability of the hydrogen bond it forms. A Tc value between 0 and –4.6 ppb/K is typically interpreted as evidence of involvement in a stable hydrogen bond, or protection from bulk water, which is associated to limited mobility of the NH. In contrast, a Tc value more negative than –4.6 ppb/K results from solvent exposure or dynamic behavior (disorder) and/or the absence of intramolecular hydrogen bonding (16, 28, 34 and references therein).

As shown in Fig. 4, most residues within the disordered N-terminal tail (residues 90-120) exhibit Tc values more negative than -4.6 ppb/K. The exception is the ∼H96-P102 segment, which may adopt some transient structure. Note that some residues in this cluster also exhibited abnormal ^15^N NOE values (*vide supra*). Residues from A113 to E146, which include the remainder of the disordered tail, the ∼V121-G126 transition coil (as identified with ^15^N NOE and R_2_/R_1_ measurements), β1, the coil connecting β1 and α1, and the N-terminal portion of α1, form an uninterrupted segment, ∼A113-E146, with strongly negative Tc values (Fig. 4), consistent with weak intramolecular hydrogen bonds. This can be interpreted as a propensity to unfold of those residues within the segment (V121-E146) that are part of the FD.

**Fig. 4.**
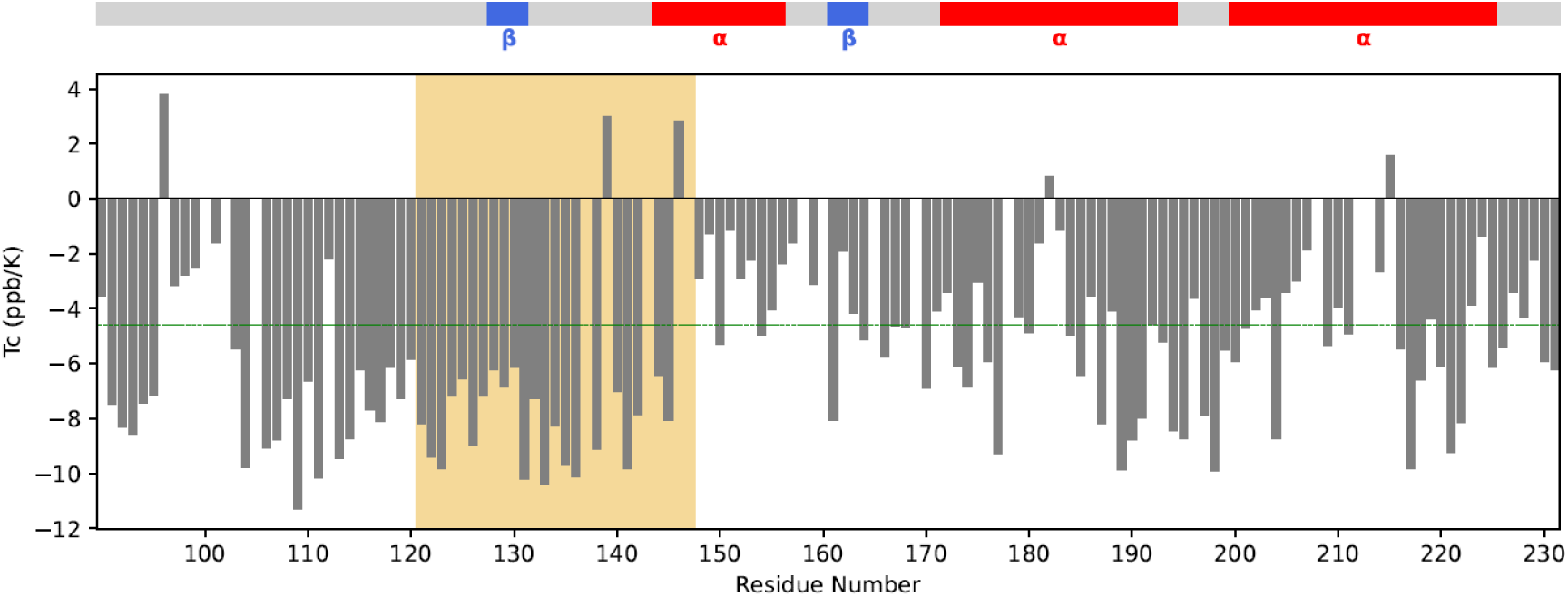
Temperature coefficients Tc of backbone amide protons. The bar plot shows the temperature coefficients TC (ppb/K) of amide proton (HN) signals in the 15-45 °C range represented along the primary sequence. The Tc values of residues giving bad linear fits have been omitted. Signals with values Tc > – 4.6 ppb/K are considered protected from the solvent or involved in permanent hydrogen bond and *vice versa*. The region shadowed in orange represents a continuous stretch of residues with Tc < –4.6 ppb/K values contiguous to the flexible 90-120 tail (see main text). The secondary structure elements are shown on the top.

The remainder of α1, and the short coil connecting it to β2, display Tc values more positive than -4.6 ppb/K, indicating greater stability. Residues in β2 and in the loop connecting β2 to α2, the so-called “rigid loop”, a misnomer given that its peculiar NMR properties are explained by alternative conformations (12, 35), show a mix of high and low Tc values, suggesting structural complexity. The “rigid loop” links not only β2 and α2, but also the two “clam-like” FD subdomains: β1-α1-β2 and α2-α3 (Figs. 5,6).

**Fig. 5:**
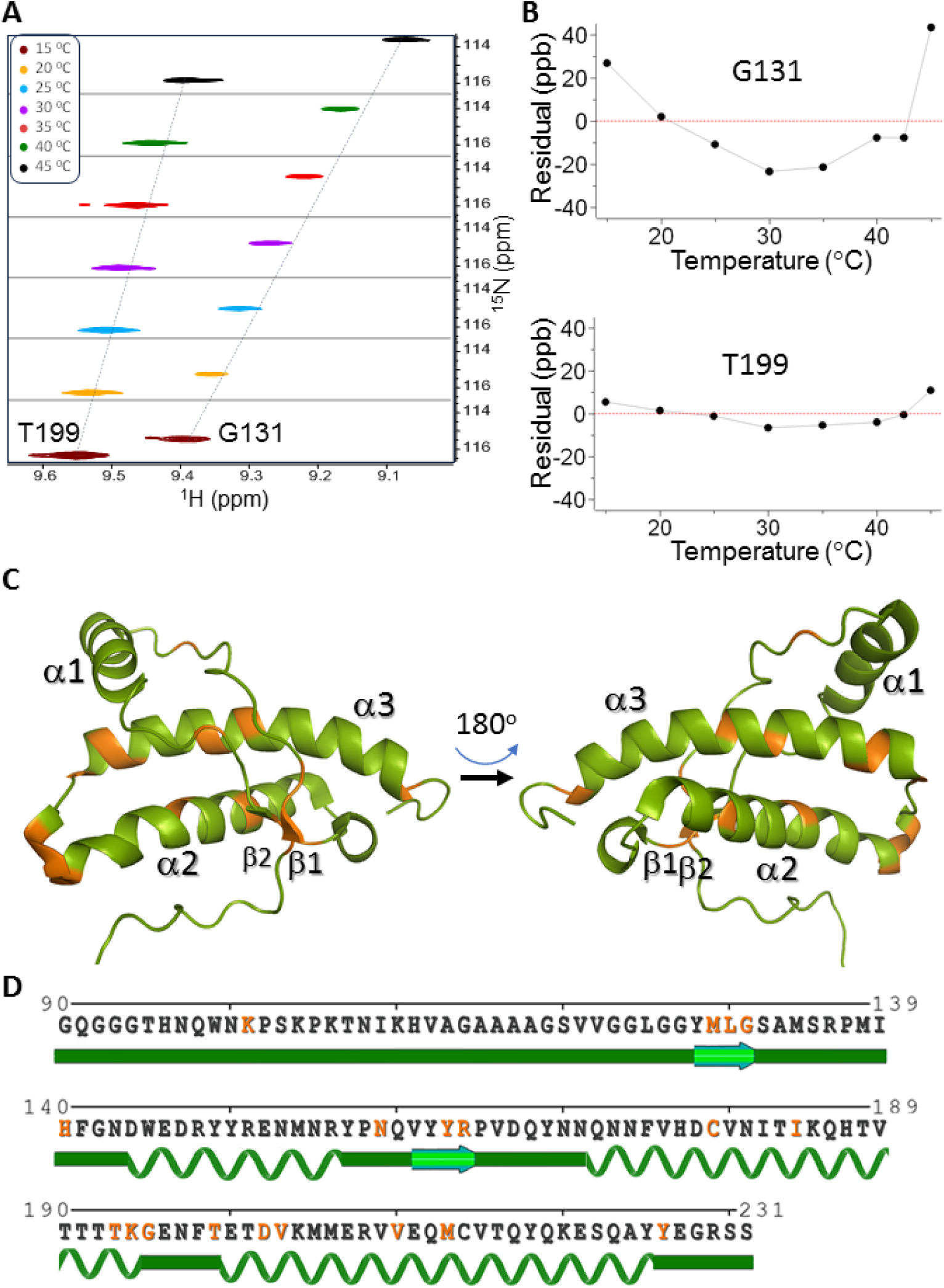
Thermally induced partial unfolding of rec(^13^C,^15^N)-BVPrP^C^90-231 tracked by solution NMR. Deviation from linearity of temperature dependent HN and N chemical shifts reports on conformational change. *(A)* Detail of stacked signals of Gly131 (non-linear) and Thr199 (linear) in N-HSQC spectra measured between 15 and 45 °C. The striped lines are shown to guide the eye. *(B)* Plots of their residual CCSD deviation. *(C)* Cartoon showing residues with non-linear CCSD response to temperature (in orange). Note how the FD features two “clam-like” halves corresponding to β1-α1-β2 and α2-α3 joined by the P165-N171 “hinge”. (D) The same information on the BVPrP^C^(90-231 sequence; residues with non-linear CCSD in orange.

**Fig. 6:**
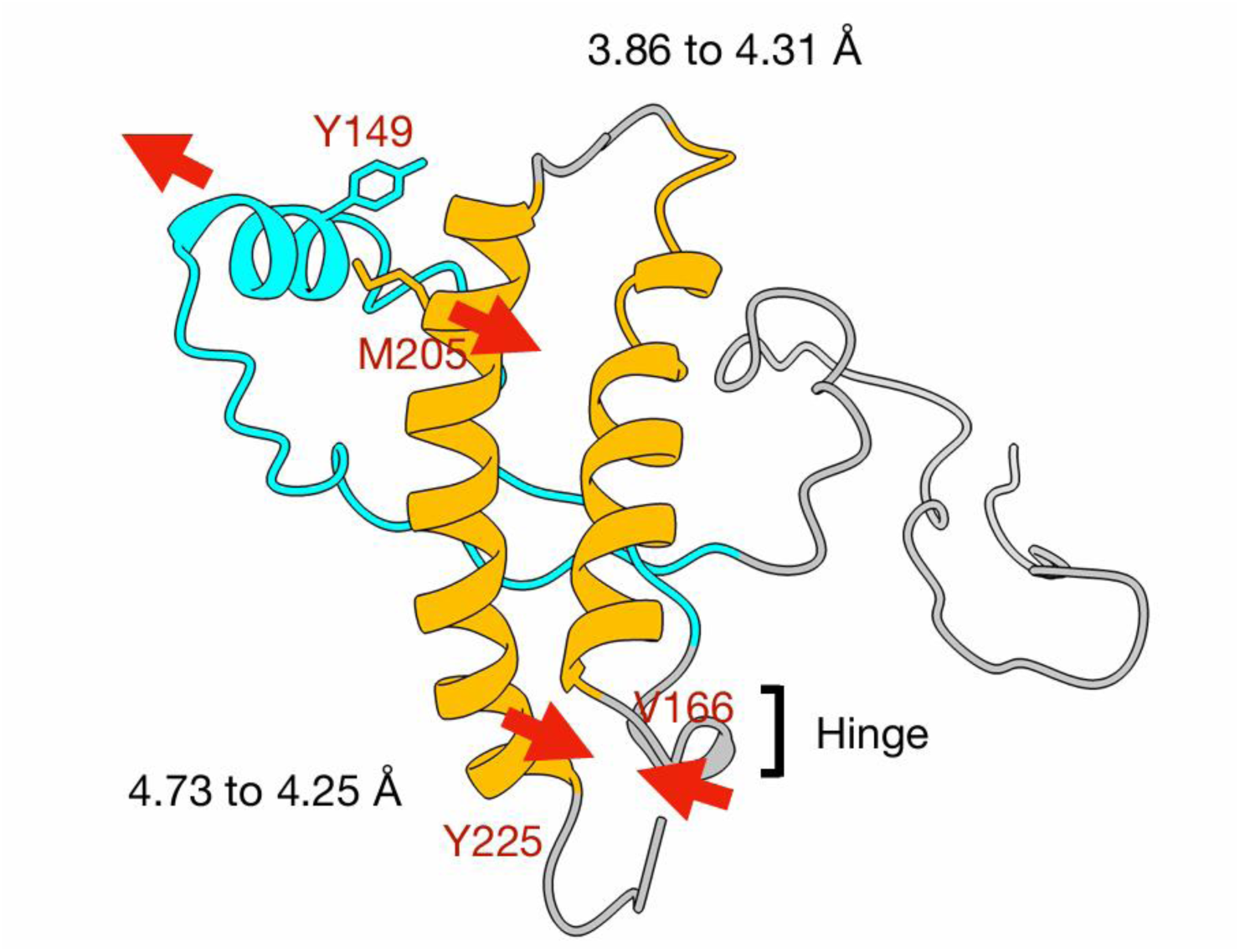
A plausible interpretation of some internal motions in BVPrP^C^ 90-231 based on the evolution of some inter-residual ^1^H-^1^H NOE distances upon heating from 25 to 45 ° C. The β1-α1-β2 and α2-α3 subdomains are colored cyan and orange, respectively; they are roughly like the two shells of a clam, joined together by the P165-N171 “hinge”, colored grey. Red arrows signal motion directions.

Both α2 and α3 display a pattern of Tc values less negative than -4.6 ppb/K in their central regions (*i.e*., relative rigidity), and more negative toward their N- and C-termini, indicating greater relative flexibility of these regions. Finally, the G195-T199 coil connecting α2 and α3 exhibits highly negative Tc values, suggesting pronounced disorder. Some specific residues display positive (Tc > 0), a rare occurrence suggesting temperature-induced conformational change (34, 36). This applies to H96 (disordered tail), I139 (β1-α1 coil), E146 (α1), I182 (α2) and V215 (α3).

Finally, we analyzed the temperature-dependent (VT) changes in amide ^1^H and ^15^N chemical shifts using N-HSQC spectra (Suppl. Figs. S4, S7). As a protein is heated, increased thermal motion generally accelerates conformational averaging and the protein is more amenable to overcome conformational energy barriers. This heating also increases the exchange rate of amide protons with water protons, leading to upfield shifts in both ^1^H and ^15^N chemical shifts (28) that are more remarkable for those residues not involved in hydrogen bond. These upfield changes are typically linear with respect to temperature. However, deviations from linearity are characteristic of residues that sample alternative, higher-energy, and less-populated conformations within the temperature range examined (34, 37).

For each peak, the ^1^H and ^15^N chemical shift drift at a given temperature was combined into a single value called the Combined Chemical Shift Difference (CCSD; Eq. 1) across the 15-45 °C range. In principle, CCSD is more sensitive to the local environment of the amide backbone (*e.g*., hydrogen bonding and solvent exposure) while offering improved noise reduction and structural robustness compared to the independent analysis of HN or N chemical shift temperature drifts. The temperature dependence of each CCSD was evaluated by fitting the data to a linear regression model (Eq. 2). This assessment of linearity is relevant for our objectives as it correlates with the conformational stability of the residue (38, 39). Residuals, representing deviations from linearity, were then plotted against temperature (Suppl. Fig. S7). Residues showing clear non-linear behavior were identified by visual inspection of these plots, and quantitatively by linearization of the residuals and calculation of their curvature.

Most residues in the folded domain (FD) showed a linear dependence of CCSD with temperature, which indicates conformational stability. However, significant deviations (non-linear CCSD residuals) were observed in residue clusters located within β1 (M129, L130, G131) and β2 (Y163, R164). Also, in the C-terminus of α2 (T193, K194 and G195, this last one located outside of α2, at the beginning of the loop connecting it with α3), and the N-terminus of α3 (D202, V203). Together with T199, also showing non-linear CCSD dependence, the segment encompassing the α2-α3 loop and immediately adjacent portions of α2 and α3 emerges a as a region exhibiting relative flexibility (Suppl. Fig. S7, Fig. 5).

There are also some other residues with non-linear CCSDs such as H140 (β1-α1 coil), N159 (α1-β2 coil), C179 and I184 (mid α2), V210 and M213 (mid α3) and Y226 (bordering the C-terminus of α3) (Fig. 5, Suppl. Fig. S7). Some additional residues exhibit CCSD *vs*. temperature plots that are modestly non-linear, and will not be considered to draw solid conclusions, except perhaps V121, V122, L125 that are more noteworthy given their clustering (Fig. 5, Suppl. Fig. S7).

Only one residue in the very high mobility 90-120 disordered tail, K101, features a non-linear CCSD plot. This can be ascribed to the fact that residues that explore a large number of conformational states with rapid interconversion typically display a linear CCSD-temperature relationship, as solvent interactions dominate (40). This trend is also apparent in the disordered 227-231 C-terminal tail, where no non-linear residues were detected.

Overall, when the relaxation data are considered together with Tc coefficients and CCSD linearity results, a coherent structural/dynamic picture emerges for BVPrP^C^(90–231). The boundary between the disordered domain and the FD, as in other PrP^C^ proteins (15, 33), is not abrupt. Instead, there is a transition region spanning residues ∼V121-G126 that is characterized by very high mobility, intermediate between that of the disordered 90-120 tail and that of a truly folded domain. Such disorder gradually decreases from its N- to its C-terminus. This is reflected in a gradient of ^15^N NOE values shifting from negative to increasingly positive, together with strongly negative Tc values. Adjacent to this segment, the ∼G127-W146 sequence also displays mobility, albeit not as pronounced as that of V121-G126. Here, positive ^15^N NOE values characteristic of a truly folded domain are accompanied by strongly negative Tc coefficients, suggesting relatively weak intramolecular HN bonds. Within this segment, β1 stands out as particularly unstable, as indicated by strongly negative Tc coefficients, CCSD non-linearity, and an unusually high R₂/R₁ ratio for G131.

The β1-α1-β2 subdomain as a whole exhibits greater mobility than the α2-α3 subdomain. Nevertheless, the coil connecting α1 and α2, as well as the C-terminal regions of both helices, show moderate mobility based on Tc coefficients and CCSD linearity. In addition, CCSD linearity analysis reveals flexibility in β1, β2 and the α2-α3 coil together with its adjacent areas: the C-terminus of α2 and the N-terminus of α3.

### Analysis of changes in long-range amide proton NOE intensity between 25 and 45°C

We next analyzed long-range ^1^H-^1^H NOEs at 25 and 45 °C. Given the lower sensitivity of the NOESY experiment, a higher protein concentration (6 mg /ml) was used. Despite this higher concentration, the behavior of this sample upon heating from 15 to 45 °C was similar to that of the 1.4 mg/ml samples described in the previous section, *i.e*., HSQC spectra did not exhibit changes other than the expected chemical shift drift (Suppl. Fig. S8). Despite the increased concentration, only a limited number of NOE cross-peaks were obtained, in stark contrast to the large number reported by Christen *et al*., who used approximately three times the protein concentration employed by us (12). This limitation is attributed to the differences of sensitivity of our experimental setup which did not include a cryoprobe and to a lower extent by the accelerated Non-Uniform Sampling (NUS) factors of 50% or 40% used, respectively, to acquire our ^15^N- and ^13^C-edited NOESY spectra (41). Despite this limitation, we were able to use the obtained NOE cross-peaks as probes to gauge the distance evolution between protein structural elements throughout the 25-45 °C heating process.

As shown in Suppl. Table I, a comparison of inter-proton distances at the two temperatures revealed that 14 of 20 decreased by approximately 0.2-0.5 Å, while two increased by a similar amount and four remained essentially unchanged. These modest variations are consistent with the small global motions inferred from the limited breadth and full reversibility of changes observed in the HSQC spectra over this temperature range (Fig. 2 and Suppl. Fig. S5).

Particularly notable are the shortened distances between the backbone amide protons of M129 and Y163, and between G131 and V161, supporting mobility within the β1/β2 ensemble (Suppl. Table I). In contrast, the distance between the amide proton of W145 and the HD proton of R148 increased substantially by 0.7 Å. This may reflect partial, reversible unraveling of α1, where both residues reside. However, movements of the R148 side chain alone cannot be excluded. The other distance that increased between 25 and 45 °C is that between the amide proton of Y149 and the HE proton of M205. These residues form part of the contact surface between α1 and α3, and the increased separation could indicate the onset of separation between the β1-α1-β2 and α2–α3 subdomains (Fig. 6), although rotation of the M205 side chain cannot be excluded.

A motion involving the P165–N171 inter-subdomain hinge might also explain the shortening of the distance between V166 and the HD proton of Y225. Similarly, the reduced distance between M154 and Y157 suggests a possible lateral displacement of α1 from the α1-β2 coil. Around their hinge. The shortening observed between Q160 and the HE proton of M213, located in the α1-β2 coil and α3, respectively, might seem inconsistent with a separation of the β1-α1-β2 and α2-α3 subdomains. However, such a change could be reconciled by a combination of twisting and partial separation of α1 from the α1-β2 coil.

Taken together, these data point to motions within the β1/β2 assembly. They do not show clear separation of the β1-α1-β2 and α2-α3 subdomains, not expected in this temperature range characterized by reversible changes and motions (Suppl. Fig. S5), but they do reveal tensions and dynamic rearrangements affecting the contact interface between them, as well as the interface between α1 and the remainder of the folded domain.

### Further heating of BVPrP^C^(90–231) results in additional unfolding and eventual collapse of partially unfolded PrP^C^ subunits to form a soluble oligomer

As mentioned above, many signals in the N-HSQC spectrum began to broaden irreversibly beyond approximately 50 °C (Fig. 2), becoming less intense and eventually completely vanishing. At 70 °C, the spectrum contained only a fraction of the peaks observed in the 15-45 °C temperature range. Although the referred loss of intensity of the signals may be due in part to the increase of exchange of the amide protons with the solvent, when the samples were cooled back to 25 °C, some of these changes persisted, and the original 25 °C spectrum was not recovered, indicating that these changes were irreversible (Suppl. Fig. S5). However, at this lower temperature, most peaks sharpened, likely due to reduced HN exchange with the solvent. For brevity, we refer to these spectra as “post-X °C” spectra, where X denotes the highest temperature to which the sample was heated. Our focus was on the post-70 °C spectrum, which features substantially fewer peaks compared to the original unheated (25 °C) sample spectrum (Fig. 7A). These peaks fall into three categories: (a) a majority corresponding to residues in the N-terminal ∼90-121 and C-terminal ∼227-231 unfolded regions; (b) peaks from residues within the folded domain (FD) with unchanged chemical shifts; and (c) a very small number of new peaks (between two and four) not present in the original 25 °C spectrum (Fig. 7A). This unique combination led us to suspect that soluble oligomers formed upon heating and cooling.

**Fig. 7:**
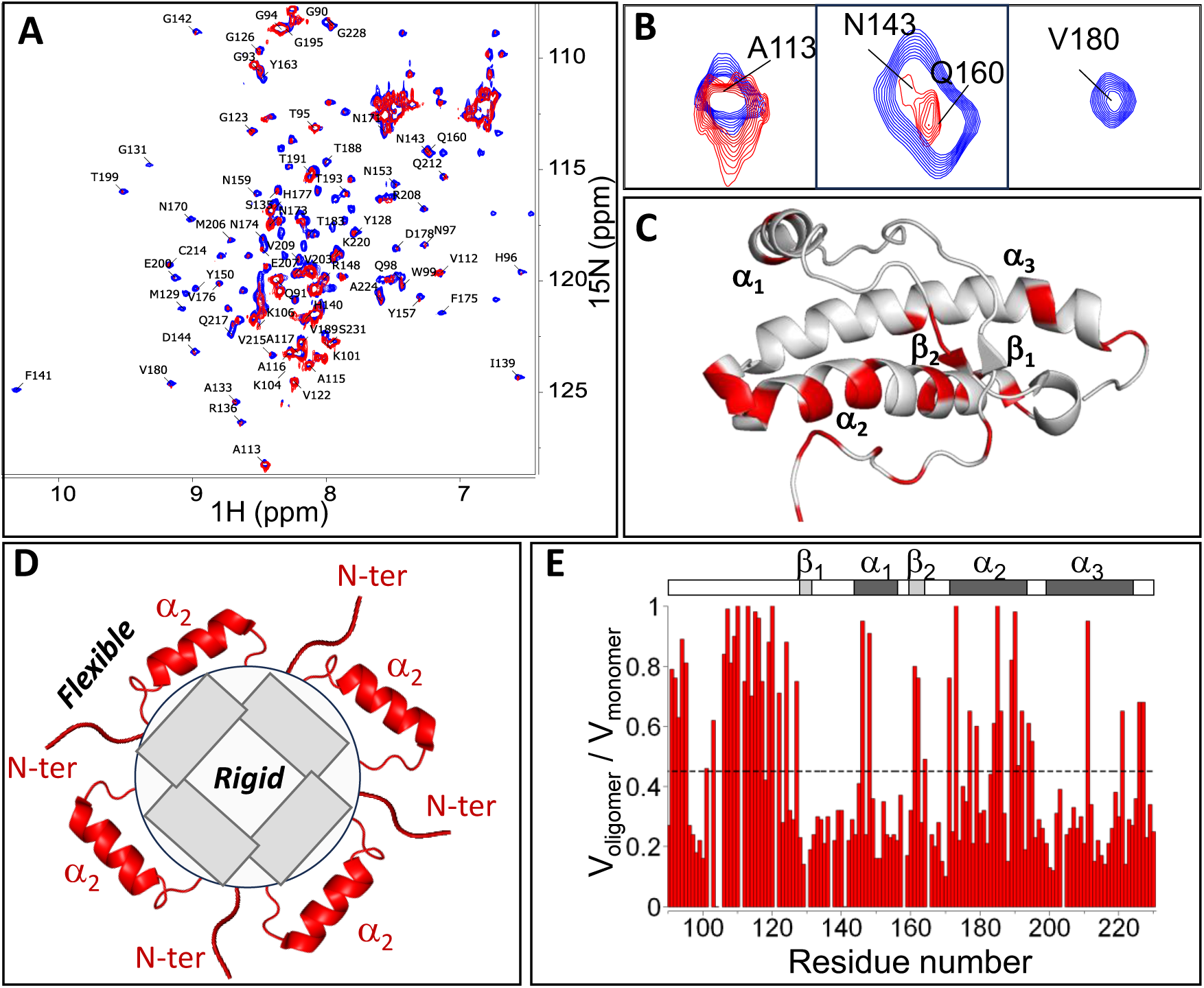
Irreversible changes upon unfolding of BVPrP^C^(90–231) above 50 °C. As thermally induced unfolding progresses, BVPrP^C^(90–231) collapses to a stable, soluble oligomeric form in which most NMR signals vanish. *(A)* Superimposed N-HSQC spectra of the initial sample at 25 °C (blue), and the same sample after being heated at 70 °C and cooled back to 25 °C (red). (*B)* Enlargement of three regions with examples of residues whose signals are still detected (A113, N143/Q160) or have disappeared (V180) and therefore must be located in substructures partially detached from the oligomer core so as to maintain a partially independent motion regime and unchanged chemical shifts. *(C)* Map of residues with HN signals detectable in the N-HSQC after heating and cooling. (*D)* Cartoon showing an oligomer with “dangling” portions that maintain independent motions and are therefore detectable in the N-HSQC spectrum. (*E)* Normalized volume ratio of signals seen in *(A)*. The stripped line at 0.45 represents the mean value of all the residues.

To investigate this possibility, we acquired DOSY spectra at 25 °C for fresh and post-70 °C samples (Suppl. Fig. S9) and calculated self-diffusion coefficients (D) using Eq. 5. The D for the fresh 25 °C sample was 0.96 ± 0.04 × 10⁻¹⁰ m²/s, whereas for the post-70 °C sample it decreased to 0.63 × 10⁻¹⁰ m²/s, strongly suggesting the presence of oligomers diffusing more slowly due to their larger size (Suppl. Fig. S9C). This latter D value likely represents a weighted average of monomeric and oligomeric species in the post-70 °C sample (*vide infra*). Using Eq. 6, the estimated molecular mass (M) from D for the fresh sample is 15 ± 2 kDa, in excellent agreement with the 17.02 kDa mass of the monomeric protein.

Next, we performed size exclusion chromatography (SEC) on post-50 °C, post-60 °C, and post-70 °C samples. The post-50 °C chromatogram shows a single peak corresponding to the BVPrP(90–231) monomer (Fig. 8). In contrast, the post-60 °C chromatogram displays a second peak, corresponding to an earlier-eluting (hence larger in size) species, with an elution time consistent with a ∼12-mer based on the column calibration. In the post-70 °C chromatogram, this peak is also present, with its relative area increasing at the expense of the monomer peak, reaching an oligomer-to-monomer ratio of approximately 2:3.

**Fig. 8:**
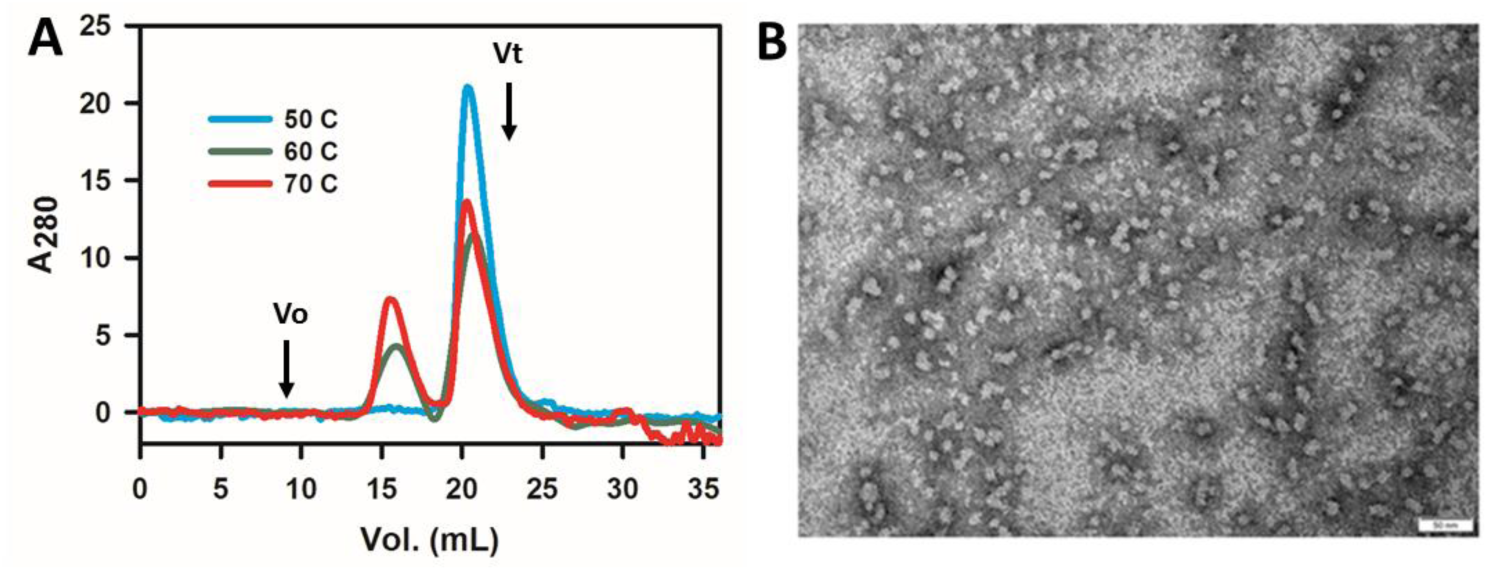
Partially unfolded BVPrP^C^(90–231) collapses into soluble oligomers at > 50 °C. *(A)* SEC analysis. (*B)* Negative stain TEM. Scale bar = 50 nm.

Going back to the DOSY results of the post-70 °C sample, we considered this monomer-to-oligomer ratio and fitted the DOSY intensity data to a double-exponential model (42) (Eq. 7) to calculate the independent Dₐ and D_b_ diffusion coefficients of the monomer and oligomer, respectively. Dₐ was fixed at 0.96 ± 0.04 × 10⁻¹⁰ m²/s, and weighing factors were used considering the 2:3 abundance ratio derived from SEC (Suppl. Fig. S9D). This yielded a D_b_ value of 0.32 ± 0.03 × 10⁻¹⁰ m²/s for the oligomeric species, corresponding to a molecular mass of 280 ± 60 kDa, or approximately 16 monomeric units, a reasonable fit with respect to the oligomer size calculated from SEC calibration (∼12 monomeric units).

Finally, negative stain Transmission Electron Microscopy (TEM) of 25 °C, post-50 °C, post-60 °C, and post-70 °C samples revealed numerous spherical particles (∼10-15 nm diameter) in the post-60 °C and post-70 °C samples, but not in the 25 °C or post-50 °C samples, providing visual confirmation of the oligomers (Fig. 8B).

### Off-track soluble oligomers report on an advanced, unstable conformer emerging in a second, more advanced phase of the unfolding process

The N-HSQC spectrum of the BVPrP(90–231) sample heated to 70 °C for 1 hour and then measured at 25 °C shows only 46 peaks with appreciable intensity, while the remaining 95 peaks either disappear or show significantly reduced intensity (Fig. 7A). Most of the visible peaks correspond to residues located in the intrinsically disordered N-terminal (∼90-120) and short C-terminal (∼227-231) sequences. However, several signals from residues within the FD remain detectable in the oligomeric sample; these must belong to structural elements that do not contribute to the oligomer core, retain their original fold (as indicated by unchanged chemical shifts), and maintain a degree of local flexibility independent of the oligomer’s overall slow tumbling (Fig. 7D).

The majority of these residues (Q172, N174, D178, V180, N181, Q186, H187, T190, T191, and T193) are located in helix α2, comprising 43% of its sequence. Residues G195 and E196, situated at the C-terminal edge of α2 in the short coil connecting to α3, are also detected. In contrast, only two residues from α3, D202 and E222, appear prominently in the spectrum. Similarly, two residues from α1 (D147 and R148, representing 15% of that helix) and two from β2 (V161 and Y162, 50% of its sequence) are observed with high intensity (Fig. 7).

Overall, substantial portions of α2 and β2 are detected in the post-70 °C oligomer-rich sample. This indicates that parts of α2 and β2 are preserved in the partially unfolded monomers comprising the soluble oligomer. These structural elements likely extend outward from the oligomeric assembly, allowing them to tumble more rapidly relative to the oligomer core (Fig. 7D). While these oligomers are artifactual, the information they afford (preservation of a substantial portion of α2 and β2 at a more advance stage of the unfolding process) is of great relevance to our purposes (see Discussion).

### Molecular Dynamics (MD) analysis

Our experimental observations consistently highlight the instability of β1 and its adjacent coil, particularly within the V121-E146 segment, which emerges as the principal hotspot for early unfolding. NOE analysis further revealed expansion of the β1-α1-β2 domain (43, 44), underscoring its intrinsic flexibility relative to the more stable α2-α3 subdomain. Based on these findings, we hypothesized that partial unfolding of β1 and its flanking coil facilitates domain separation and may provide the structural plasticity required for PrP^C^ to undergo fit-induced adjustment at the PrP^Sc^-PrP^C^ interface during the initial stages of conversion. Therefore, we used plain MD simulations as an alternative and complementary approach to probe PrP^Sc^-assisted unfolding of BVPrP^C^(90–231). Unlike solution NMR, this method allows exploration of an ensemble where the disordered N-terminal tail, up to residue A120, is attached to a PrP^Sc^ templating surface and adopts the PrP^Sc^ conformation. To model this, we performed MD simulations of BVPrP^C^(93–231) with its 93-120 tail attached to a stacked tetramer of PrP^Sc^, serving as a surrogate for a longer PrP^Sc^ stack. Since no BVPrP^Sc^ structure has yet been resolved, we used RML MoPrP^Sc^ (PDB: 7td6, ref. 5) as an approximate model of BVPrP^Sc^. Accordingly, the BV^C^(121–231) FD was attached to the C-terminus of an RML MoPrP^Sc^ tetramer with an extra 93-120 upper rung (Fig. 9A). An initial 300 ns simulation at 310 K showed that the FD structure was maintained throughout, with the three α-helices remaining intact, although undergoing considerable vibrational motions. The extreme N-terminus of the attached tail detached early from the PrP^Sc^ surface and fluctuated throughout the simulation, detaching and reattaching (Suppl. Video S1).

**Figure 9.**
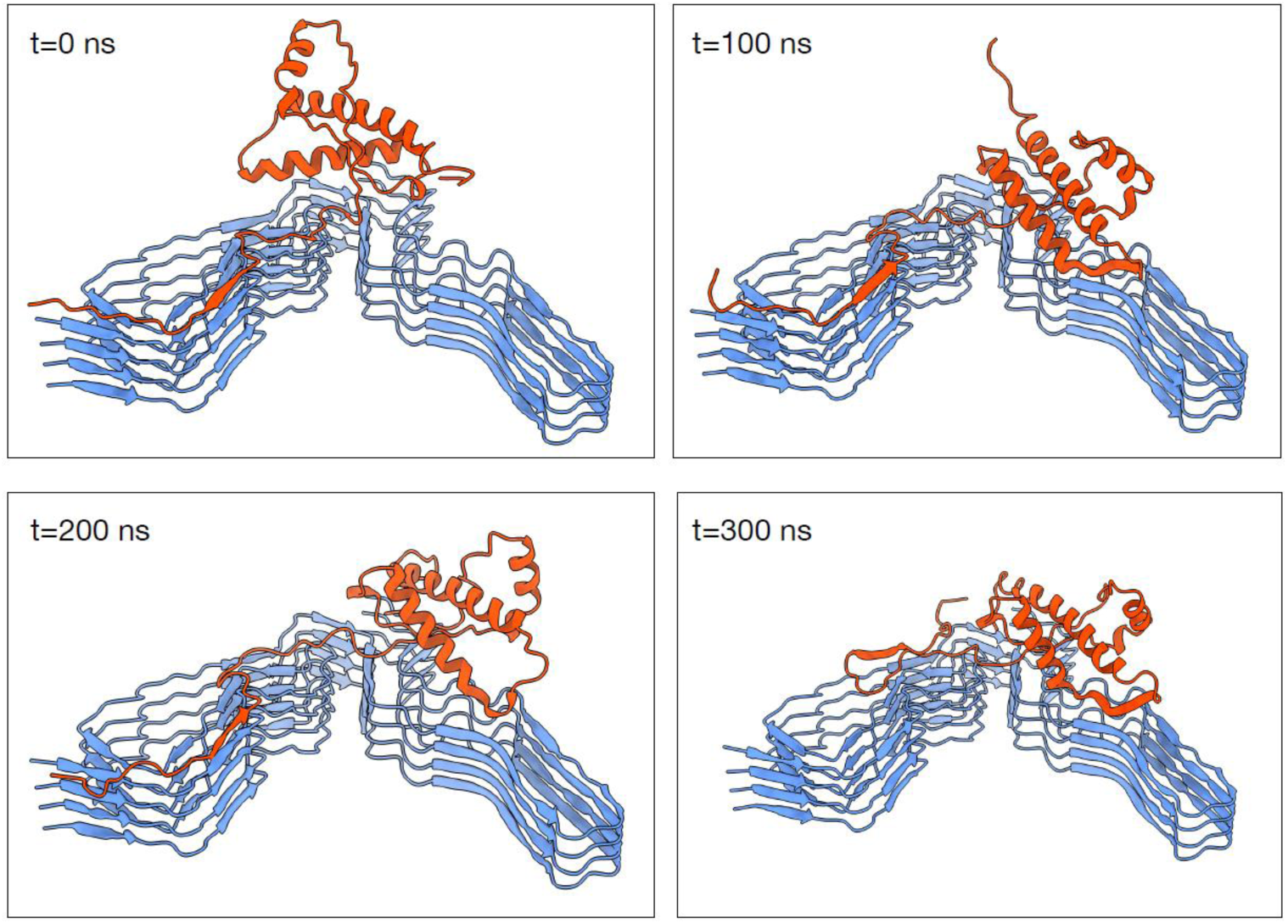
Molecular Dynamics simulation of the evolution of BVPrP^C^(93–231) with its 93-120 tail attached to PrP^Sc^ and already converted to the PrP^Sc^ conformation. Trajectories were obtained at 400 K in explicit water. *(A)* t = 0 ns; *(B)* t = 100 ns; *(C)* t = 200; *(D)* t= 300 ns. Frames taken from Suppl. Video S2.

To accelerate unfolding, the simulation was repeated at 400 K. The results, representative of three trajectories, resembled those at 310 K but displayed more intense fluctuations. However, once again, the three helices emerged unscathed from the simulation. More intense motions were also seen for the connecting coils, in particular 121-143, which underwent extensive changes of conformation. While the focus of these simulations is the FD, it is of interest that at this temperature the 93-120 coil initially attached to the PrP^Sc^ surface exhibited substantial fluctuations, with its N- and C-terminal sections detaching from the template, while its central ∼103-110 segment remained more stably attached to it (Suppl. Video S2 and Fig. 9). Quantitative RMSF analysis (Fig. 10) revealed very high fluctuations in the disordered N- and C-terminal sequences, while coils β1-α1, β2-α2, and α2-α3, along with most of α1 and the termini of α2 and α3, showed RMSF values that were also elevated but not as much. The central regions of α2 and α3 around the disulfide bridge exhibited low RMSF values, as did the β1 and β2 strands and the α1-β2 coil.

**Fig. 10.**
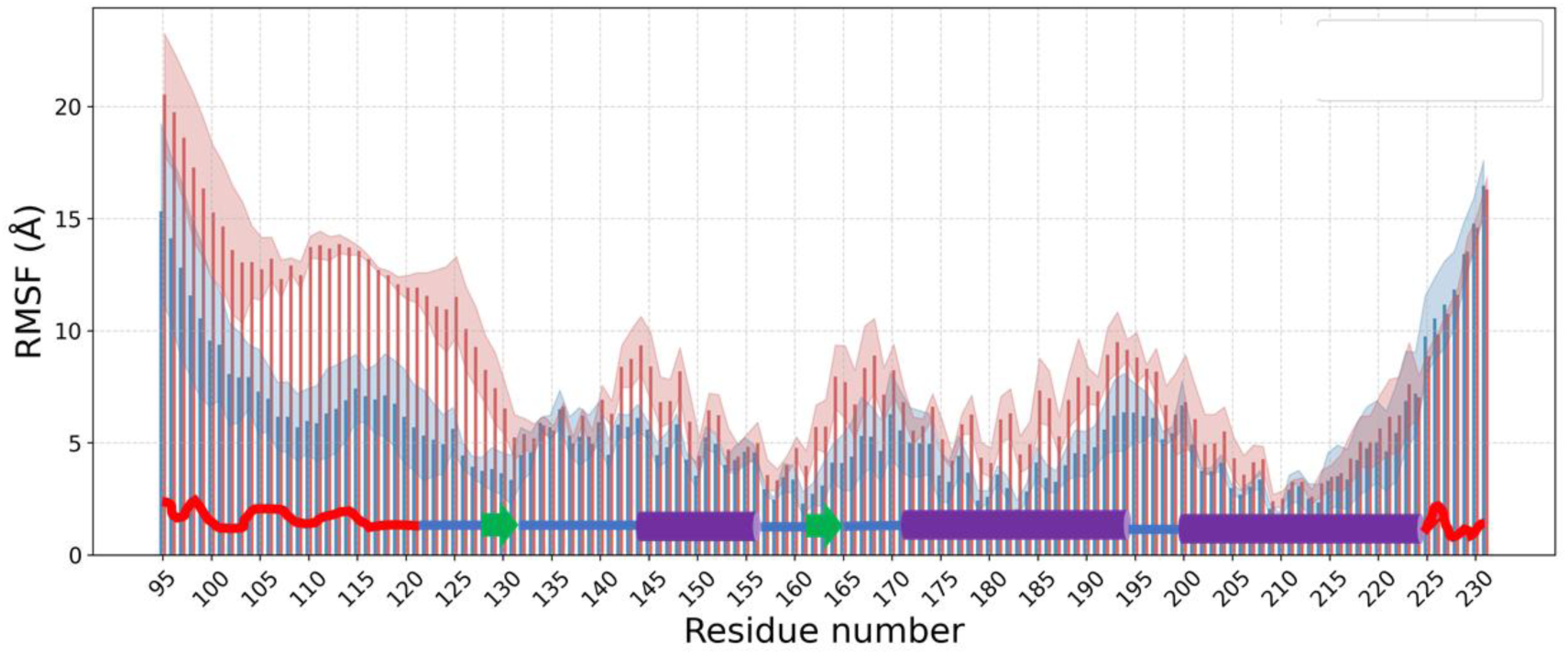
Amino acid residue RMSF throughout MD simulations at 400 K of BVPrP^C^93-231. Blue: 93-120 tail attached to a PrP^Sc^ tetramer. Red: 93-120 tail free. MD simulations are 300 ns. Bars represent means of 3 trajectories. Shaded areas cover +/- sd. Superimposed: secondary structure elements.

We also ran MD simulations of free BVPrP^C^(90–231), without the N-terminus attached to a PrP^Sc^ surrogate tetramer, at both 310 and 400 K. Some trajectories, one of two at 310 K and two of three at 400 K, resembled those of the attached system, albeit with somewhat larger fluctuations (see Suppl. Videos S3 and S4 compared to S1 and S2, and RMSF plots in Fig. 10). However, in one 310 K and one 400 K trajectory, motions were considerably more intense overall, including a drift and separation of α1 from the rest of the FD ensemble (Suppl. Videos S5, S6 and Suppl. Fig. S10, S11).

## Discussion

Since the PrP^Sc^ templating surface is relatively inert, the FD of PrP^C^ necessarily has to adapt to it through unfolding and refolding. We investigated the initial stages of FD unfolding using thermally induced unfolding coupled with solution NMR. Although a pure FD construct, *i.e.*, BVPrP^C^(121–231) might seem the most appropriate choice, we chose BVPrP^C^(90–231) to facilitate comparison with prior studies that used a variety of constructs (121-230, 113-230 or 90-230) under diverse unfolding conditions (15, 22, 46, 47). On the other hand, studying experimentally an FD tethered to PrP^Sc^ would be ideal but is technically intractable; we therefore examined BVPrP^C^(90–231) alone, assuming that FD unfolding is largely similar in both contexts.

We selected the bank vole sequence because BVPrP^C^ can adapt to all known PrP^Sc^ strain conformations (48, 49). This generic convertibility could be explained by a broad panel of conformers in the folding landscape of BVPrP^C^. Given that we aim at elucidating general mechanisms of unfolding/conversion, studying the universal conversing substrate seems the best choice.

Our results map the likely initial unfolding route of the BVPrP^C^ FD (Fig. 11). A prominent early event is disruption of the short antiparallel β-sheet (β1: Y128-G131; β2: V161-R164), supported by four observations: 1) relaxation data: G131 exhibits the highest R₂/R₁ values at both 25 °C and 45 °C, with V161, Y162, and Y163 showing also high values at 45 °C (Fig. 3), indicating significant ps-ns motions in both strands, especially at elevated temperature; 2) negative Tc values: strongly negative Tc values of β1 residues (Fig. 4) suggest weakened intra-molecular hydrogen bonding. 3) Non-linear CCSD temperature dependence in multiple residues of β1 and β2, indicating conformational sampling across 15-45 °C (Fig. 5). This is indicative of exploration of alternative conformations throughout the 15-45 °C temperature ramp; 4) NOE intensity changes: reduced NOE cross-peak intensities between β1 and β2 at 45 °C compared with 25 °C (Suppl. Table I) indicate motions affecting the β1/β2 ensemble.

**Fig. 11.**
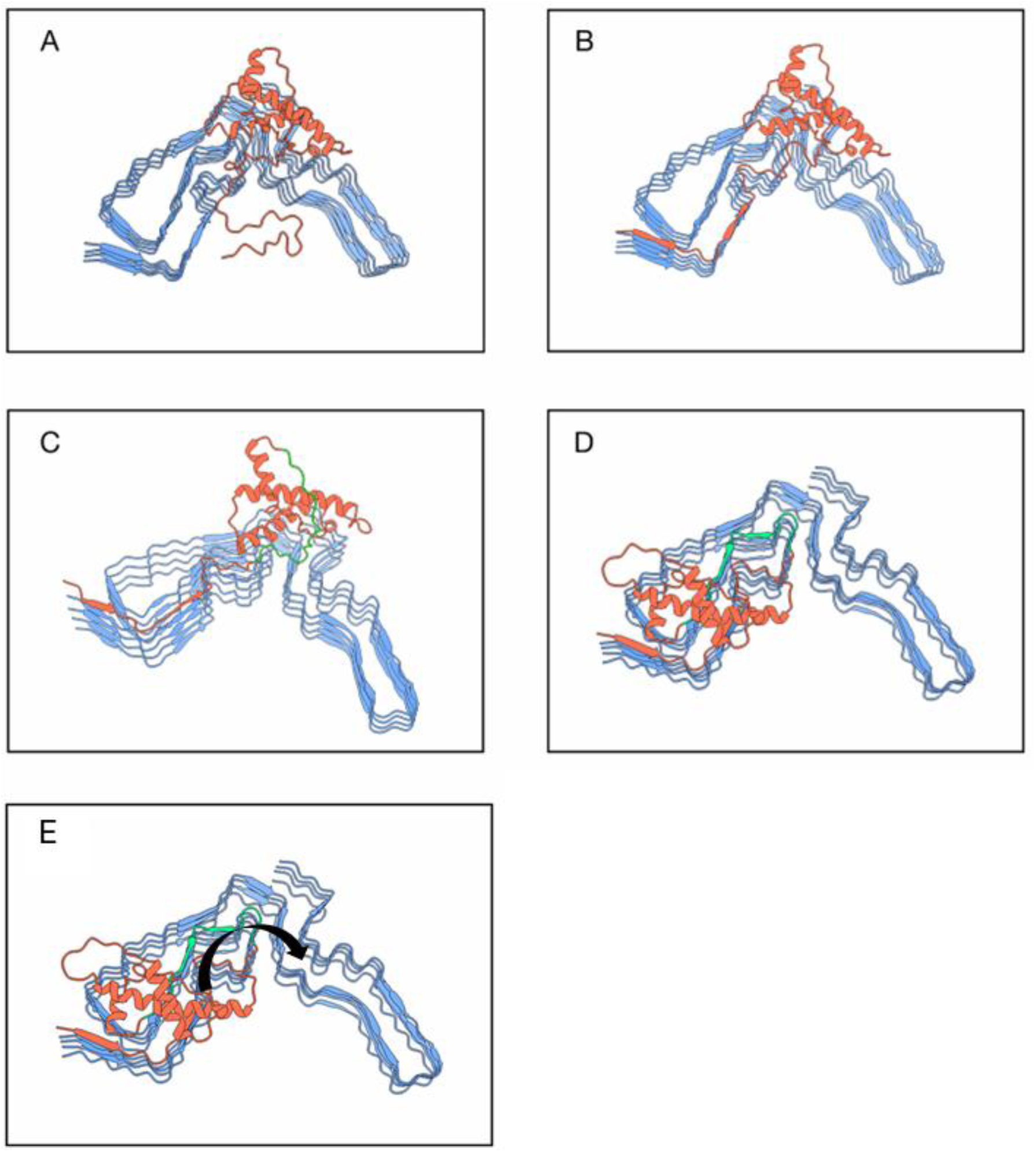
Towards an early timeline of PrP^C^ unfolding to refold to PrP^Sc^. *(A)* PrP^C^ encounters PrP^Sc^ on the cell surface. *(B)* The flexible/disordered stretch (∼90-120) of PrP^C^ is easily trapped and templated by the (∼90-120) templating surface of PrP^Sc^. *(C)* The PrP^C^ FD now hovers in front of the essentially inert PrP^Sc^ surface; while stable, some regions within the FD explore partial unfolding. The ∼V121-D144 region (green), encompassing the coil N-terminal to β1, β1, the β1-α1 coil and the N-terminal stretch of α1, constitutes the most active region. *(D)* The partially unfolded ∼V121-D144 region is the most likely to be trapped by the templating surface. However, this is speculative at this point and therefore the green color is maintained. The FD is now reduced to ∼140-231 and lacks β1. *(E)* α1 and the β2-α2-α3 ensemble drift apart around the α1-β2 coil, and eventually α2-α3 from β2 around the β2-α2 “rigid” coil to gravitate towards the C-terminal “wing” of PrP^Sc^. Given that no BVPrP^Sc^ structure has been resolved yet, the structure of RML, a mouse PrP^Sc^ (PDB 7td6) has been used, which does not affect the conclusions of this analysis.

These changes occur alongside high mobility in the coil N-terminal to β1 (Fig. 3A,B), a structural transition zone between the disordered tail and the ordered FD (12, 50). The β1–α1 coil also shows strong flexibility, with continuous negative Tc values (Fig. 4). All in all, the 121-144 region presents itself as a continuous segment of residues with higher than average flexibility.

Additional early-flexibility sites include: 1) the segment spanning the C-terminus of α2, the α2-α3 coil, and the N-terminus of α3; 2) the C-terminus of α3 and 3) the region adjacent to β2 (α1-β2 coil, β2, β2-α2 coil). These are shorter and less prominent than the 121-144 region, as judged by Tc magnitude and extension. Consider for example the 187-201 segment around the α2-α3 coil, a block of 15 residues with highly negative Tc values, compared to 24 in the 121-144 segment (Fig. 4). Or the even shorter C-terminal stretch of α3 (216–225). This, together with other considerations to be discussed below, signal the121-144 region as a strong candidate to initiate the unfolding of the FD following attachment of the 93-120 flexible coil to PrP^Sc^, in a zipper-like fashion.

Our data align broadly with prior PrP^C^ *misfolding* (see the introduction for the difference we make between *misfolding* and unfolding) studies. Russo *et al.* identified β1, β2, α1, and the α2-α3 coil as part of an “excited state” in HuPrP(90–231), with α2 and α3 largely preserved (15). However, unlike BVPrP^C^(90–231), HuPrP^C^(90–230) (but not HuPrP(23–231)), formed a thermal unfolding intermediate at 61 °C absent in our experiments (Fig. 1). Such partially unfolded forms (PUFs) occur in some, but not all, PrP sequences under specific unfolding conditions (18, 45, 51–53). Russo *et al*. also observed that HuPrP^C^(90–230) underwent complete unfolding in a fully reversible way, whereas in our experiments BVPrP^C^(90–231) underwent irreversible unfolding above ∼50 °C, forming soluble β-sheet-rich oligomers (15). We regard these oligomers as useful artifacts capturing some transient later stage unfolding intermediates rather than physiologically relevant on-pathway species (Suppl. Fig S12). We are interested in the unfolded conformer that exists just instants before they collapse into the oligomers, not in the oligomers *per se.* We hypothesize that in the presence of the PrP^Sc^ template, such unfolded conformers, tethered as they are to PrP^Sc^ through their N-termini, would collapse onto the templating surface, rather than onto similar partially unfolded PrP^C^ companions to form oligomers (Suppl. Fig. S12).

Having clarified this, what do these artifactual oligomers obtained at >50 °C in our thermal unfolding tell us? Their analysis indicates: 1) their immediate precursors form after earlier structural changes, likely involving the ∼121-144 segment, as discussed above; 2) Substantial portions of α2 and β2 persist within them; and 3) they exhibit decreased α-helical and increased β-sheet content (see comparative CD spectra in Fig. 1). The presence or absence of α3 in these intermediates remains uncertain. Absence of more signals from α3 might be the result of NMR signal broadening if α3 is also preserved to some extent but is more intimately associated to the slow-tumbling oligomer core than α2 (Fig. 7).

Similar oligomers, typically generated at acidic pH with denaturants or salts, share high solubility, β-sheet enrichment, and concurrent oligomerization/secondary-structure changes (19, 22, 54–63). Their generation by thermal unfolding alone has also been described (43, 64–66). The virtually superimposable kinetic traces of increase in β-sheet content and oligomerization support the notion that these two changes occur in a nearly simultaneous fashion (56). This lends support to the notion that the oligomers trap a fleeting monomeric PrP unfolding conformer. Overall, the literature suggests that such intermediate preserves much of α2-α3 while β1-α1-β2 partially refolds into β-sheet structure. Thus, Honda *et al*. showed, using deuterium exchange coupled with NMR analysis, that protection factor across the β1-α1-β2 subdomain is orders of magnitude lower than that of the α2-α3 subdomain immediately prior to oligomerization. This strongly suggests generalized unfolding of β1-α1-β2, with some α1 and coil segments eventually converting to β-sheet and collapsing into the oligomeric form (19).

The atomistic structure of the oligomers has not been deciphered to date. However, experimental data suggest that it might involve separation of these subdomains and domain-swapping-like interactions. Serpa *et al*. used a combination of mass spectrometry combined with chemical crosslinking, hydrogen/deuterium exchange, limited proteolysis, and surface modification to probe the conformation of PrP oligomers. These studies provided a large number of restraints that were fitted to a structural model in which β1-α1-β2 has unfolded, flattened, increased its β sheet content and drifted away from the α2-α3 subdomain to interact instead with the modified α1-α2 subdomain of a contiguous subunit of the oligomer, in an arrangement that is reminiscent of a “runaway” domain coupling (62).

While oligomers are very soluble, and therefore amenable to solution NMR analysis, studies had consistently shown disappointingly uninformative spectra in which only signals arising from the long N-terminal and short C-terminal intrinsically disordered segments (∼23/90-120 and ∼225-230, respectively) are detected (22, 59, 60). This is a consequence attributable to severe line broadening resulting from slow tumbling of the large oligomer in solution, together with partially independent fast motions of the dangling flexible “tails” (18, 59). This suggests that our oligomers have some distinct architectural characteristics, namely, α2 is substantially preserved, and it must constitute an ensemble that likely juts out of the oligomeric core (Fig. 7).

In this respect, our results do align with data reported recently by Russo *et al*. As mentioned earlier, when these authors incubated HuPrP^C^(90–230) during 1 h in a temperature range of 15-80 °C, its unfolding was fully reversible. However, when they heated it for several hours at 61 °C they obtained a certain amount of irreversible oligomers, albeit with an apparently lower yield than ours with BVPrP^C^(90–231) (15). This different behavior might result from intrinsically different unfolding properties of BVPrP and HuPrP. When they compared the ^1^H-^15^N HSQC spectra of this “post-61°C” sample and the original one, both at 25 °C (an experiment essentially similar ours as reported in Fig. 7) they observed, like us, signal loss and broadening for a large number of residues in the FD without significant chemical shift changes. But similarly to our results, some residues located within the FD were less affected, namely most residues in α1, the C-terminus of α2. Except for residues in β2, that were severely suppressed, there is a relatively good coincidence with our results. The mixture of oligomers and monomers (∼40:60 in our case, likely fewer oligomers in their case) adds complication to the interpretation of data. Isolation and characterization of the oligomers alone, ongoing, should help. Finally, it is noteworthy that the oligomers obtained and described by Russo *et al*. were shown to be “on-pathway” towards generation of a HuPrP(90–230) amyloid, underscoring the differences between studying *misfolding* and *unfolding* in the context of PrP^Sc^-templated propagation, even though some conformers (in this case those immediately preceding the oligomers) might be common.

Molecular dynamics simulations of BVPrP(90–231) with its 90-120 tail attached to a static PrP^Sc^ tetramer surface support and complement the timeline suggested by the experimental studies. The most flexible/moving parts of the FD are the coils, as shown by the RMSF plot (Fig. 10 and Suppl. videos S1, S2). In contrast, the 3 α-helices are more resilient, they stretch and “breath” significantly around their original conformation, without losing their structure. The same is true for the two β sheets, that keep on separating from one another and temporarily lose their canonic parameters, only to recover them. The relatively low RMSF of β1 interrupting a sequence of residues with high RMSF spanning from A120 to α1 is apparently at odds with our conclusion that the entire A120-D144 is the most likely hotspot for unfolding/refolding. However, it should be noted that the β1 sequence retains its β secondary structure in PrP^Sc^, where it becomes β4 (3, 5). Therefore, it is conceivable that it behaves like a small “β-sheet pre-assembly unit”, fitting into the corresponding β4 sequence *en bloc*.

While the overall behavior of BVPrP^C^(90–231) with its N-terminal tail unattached was very similar in MD simulations carried out both at 310 and 400 K to that of BVPrP^C^(90–231) with its tail attached to a PrP^Sc^ surrogate, the general mobility of all elements of its sequence was higher (Fig. 10). In two of the trajectories, separation of α1 from the rest of the FD was seen (Suppl. Fig. S10 and Suppl. videos S5-S6). This agrees with analysis of NOEs suggesting a tendency of the β1-α1-β2 and α2-α3 subdomains to separate, and more specifically, of α1 to drift apart from the rest of the FD by a torsion at the Y157-Q160 coil connecting it with β1. However, the occurrence of this event only in some of the trajectories suggests that it is rare and therefore it likely takes place at later times throughout the unfolding process. Separation of the β1-α1-β2 and α1-α2 subdomains is *sine qua non* to complete templating of the FD by the flat PrP^Sc^ surface for obvious geometrical reasons (2, 14). Evidence of such molecular event has been previously gathered by MD (17, 67–69) and experimental data using FRET (70).

Collectively, our data suggest a plausible early unfolding pathway for PrP^C^ conversion in the presence of PrP^Sc^ (Fig. 11): initial ∼90-120 tail attachment and templating; destabilization of ∼V121-D144 (the ∼121-125 segment is intrinsically extremely flexible) together with concomitant separation of β1–β2 and eventual piecemeal templating of the destabilized FD. The V121-D144 region is likely the first major FD segment to unfold and be trapped, producing a truncated ∼140-231 FD tethered to the templating surface through a small ∼138-144 coil (Fig. 11D). Such shortened FD would be analogous to the destabilized HuPrP^C^(137–230) conformer described by Hosszu *et al*. (71).

HuPrP(137–230) exhibits substantially less stability than HuPrP^C^(119–230), with a 20-fold reduction in the equilibrium between the native and unfolded states (K(N/U) ∼2000), a ∼10 °C reduction in the mid-point for thermal unfolding, and decreased protection of β2 and α1 as determined by deuterium/hydrogen exchange and a substantially increased tendency to fibrillize (71). While we cannot be certain that a hypothetical post ∼120-140-trapping FD as shown in Fig. 11D is identical to the PrP137-230 conformer described by Hosszu *et al*., whose atomic structure is not available yet (71), it is reasonable to think that they must be similar. Therefore, it is very likely that a PrP^Sc^-tethered ∼140-231 FD would exhibit a substantially increased lack of stability, with a higher propensity of specific regions to unfold and eventually be trapped and templated piecemeal by the PrP^Sc^ surface.

At some point, α2-α3 must separate from the β1-α1-β2 block, part of which will have already been converted to the PrP^Sc^ conformation and gravitate towards the C-terminal lobe of the PrP^Sc^ templating surface (Fig. 11E). MD simulations suggest that this happens at a later stage. A substantial portion of α2 and bits of α3 resists in its native conformation at high temperatures (roughly equivalent to advanced time of evolution) in our experimental unfolding paradigm. This supports the notion that the α2-α3 ensemble is structurally solid and might be the last portion of the FD to unfold and convert to PrP^Sc^. This is also in agreement with geometric considerations.

It should be emphasized that the unfolding/refolding timeline we propose certainly contains speculative elements. It represents a plausible succession of events that fit our experimental and modelling data. However, alternative timelines are possible. For example, the disordered short C-terminal Y226-S231 segment, rather than the N-terminal one might be first in attaching to the PrP^Sc^ templating surface. However, this seems less likely given the steric hindrances derived from the presence of the GPI anchor attached to S231. Furthermore, the sequence is very short and might detach (Fig. 9) before there is a chance of further extension.

Another alternative is that once the ∼90-120 tail has attached/converted, a region other than ∼121-140 is first to unfold and be trapped. It could be one of the coils, for example G195-T199 and surrounding areas: the C-terminus of α2 and the N-terminus of α3. This segment contains a number of residues with low Tc values and non-linear CCSD plots. However, the geometry of the FD and the distance to this segment to its iso-sequential “docking area” in the PrP^Sc^ surface militate against this option. Nevertheless, it cannot be excluded that more than one reaction coordinate exists, all taking place in parallel. Nonetheless, the proposed pathway integrates our experimental and modeling data and provides a structural framework for future atomistic models of PrP^Sc^ propagation.

## Materials and Methods

### Recombinant protein production

The recombinant bank vole (BV) PrP(90–231) protein, corresponding to the I109 natural polymorph (GenBank accession number PQ 327920), was expressed in bacteria and purified as described previously (72). Briefly, the Open Reading Frame of the wild-type I109 bank vole Prnp gene was obtained from genomic DNA by PCR and cloned into the pOPIN E expression vector, which adds an amino-terminal 6x-His tag to the encoded protein, without any linker. We decided to keep the N-terminal 6x-His tag used to facilitate purification for practical reasons, like others before us (19, 21, 59, 60, 62). It has been experimentally established that the His_6_ tag, at the end of the intrinsically unfolded ∼90-120 segment, does not influence unfolding of the FD (21). E. coli Rosetta™ (DE3) competent cells were transformed with the expression vector using standard molecular biology procedures. Expression of ^13^C,^15^N labelled recombinant protein was achieved by growing the bacteria in two flasks with 25 ml each of LB broth overnight and then for 4-5 h in two 2L flasks containing each 500 ml of minimal medium supplemented with 3 g/L of uniformly labelled ^13^C-glucose and 1 g/L of ^15^N-NH_4_Cl. Upon isopropyl β-d-1-thiogalactopyranoside (IPTG) induction, inclusion bodies containing the protein were harvested. Purification of the protein was performed with a histidine affinity column (HisTrap FF crude 5 ml, GE Healthcare Amersham). After elution in buffer consisting of 20 mM Tris-HCl, 500 mM NaCl, 500 mM imidazole and 2 M guanidine-HCl, pH 8, the quality and purity of protein batches was assessed by BlueSafe (NZYtech) staining after electrophoresis in SDS-PAGE gels (BioRad). Finally, guanidine-HCl was added, to a final concentration of 6 M, for long-term storage of purified protein preparations at -80 °C.

Protein folding was performed by dialysis against 10 mM sodium acetate, pH 5 using 3 kDa cutoff membrane Slide-A-lyzer dialysis cassettes (Thermo), with three changes of dialysis buffer. Aggregated material was removed by a brief centrifugation at 18,000 g in a Beckman Microfuge 22R tabletop centrifuge. Protein concentration was estimated spectrophotometrically, using an ɛ_280_ of 26000 and a MW of 17.02 kDa. The folded protein was concentrated as needed using Amicon centrifugal devices and immediately used for different analyses or kept at 4 °C for a few days in between experiments. In such case, the quality of the protein was assessed by H-N HSQC spectra.

### NMR spectroscopy

All NMR experiments were carried out in solution using a Bruker NEO 750 MHz spectrometer equipped with a PA-TXI-HFCN (^1^H-^19^F/^13^C/^15^N) room temperature probe with an automated tuning and matching system, and temperature control for VT. The spectrometer control software was TopSpin 4.3.x.

#### NMR sequence assignments

The protein, in 10 mM sodium acetate buffer, pH 5, was supplemented with D_2_O (10%). The final protein concentration was 6 mg/ml. The assignment of Ca, Cb, C, N, HN and Ha resonances at 25 °C was carried out with the following 3D standard triple resonance experiments: HNCA, intra-HNCA, HNCACO, CBCANH, intra-CBCANH, CBCA(CO)NH, HNCO, HN(CO)CA, HNHa, HNCAHA, HCCH-TOCSY, ^15^N and ^13^C edited TOCSY-HSQC and ^15^N and ^13^C edited NOESY-HSQC. The spectra were processed with TopSpin and analyzed with CCPN software v3.2 (73) guided by the chemical shifts reported by Christen *et al*. for BVPrP^C^(125–231) (12) and deposited under the BMRB accession code 15824. Additional unpublished chemical shift values at 25 °C were kindly provided by Simone Hornemann, University Hospital, Zürich. The assignment of the N-terminal residues 90-118 was also assisted by comparison with the analogue sequence of mouse Q216R-PrP (16, PDB entry 51007). The chemical shift assignment of N and HN resonances at different temperatures was carried out by tracking peak trajectories in a series of N-HSQC Spectra (*vide infra*) over the temperature range. Spectra were processed with TopSpin and the peak positions were analyzed using MestreNova software (Mestrelab Inc.) ^1^H, ^13^C and ^15^N chemical shifts were referenced indirectly to external 2,2-dimethyl-2-silapentane-5-sulfonic acid (DSS) reference. Details on these and the rest of NMR experiments performed are listed in Suppl. Table II.

### Variable Temperature (VT)-NMR data analysis

N-HSQC spectra were acquired at increasing temperatures from 15°C to 45°C in 5°C or 2.5 °C increments. Each spectrum was referenced using the acetate signal of the buffer as secondary reference, and chemical shift changes for each amide proton-nitrogen (HN) cross-peak were tracked across the temperature range. The ^1^H and ^15^N drift of a peak in the N-HSQC spectrum at a certain temperature of the series was analyzed by the Combined Chemical Shift Difference (CCSD) (74) (Eq. 1)

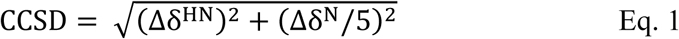

Where Δδ^HN^and Δδ^N^is, respectively, the change in ^1^H and ^15^N chemical shift respect to the reference values of the spectrum acquired at the lowest temperature of the series. The experimental CCSD for a specific amino acid at each temperature, was linearized using MS-Excel software by fitting the data to the following equation (Eq. 2):

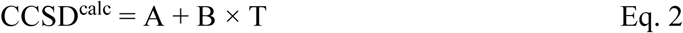

Here, CCSD^calc^ represents the calculated linearized CCSD value for each data point, T is the temperature, and A and B are the regression coefficients. The residual error (defined as CCSD - CCSD^calc^) was calculated and plotted against temperature (38, 39). Residual errors showing linear or non-linear behavior were identified by visually inspecting the plots and quantitatively by curvature calculation through second-order linearization of the residuals and evaluation of the RMSD factor. Plots of the residuals were represented by SciDAVis software (scidavis.sourceforge.net) or by a homemade python script based in the *matplotlib* library. The non-linearities identified in the CCSD study were represented color-coded and overlaid onto the primary structure sequence using Polyview 2D software (75).

### Evaluation of Temperature coefficients (T_C_)

Temperature coefficients were determined by analyzing the chemical shift of the amide proton (34 and refs. therein) of each residue in a series of N-HSQC spectra measured in the temperature range of 15 °C to 45 °C, in 5 °C or 2.5 °C increments. The HN chemical shift versus temperature for each amino acid was linearized by MS-Excel software, and the temperature coefficient (*T_C_*, in ppb/K) was taken from the slope. *T_C_* values were plotted along the PrP^C^sequence.

### NOE measurements

3D ^15^N-edited ^1^H-^1^H NOESY spectra were acquired (sequence *noesyhsqcfpf3gpsi3d)*) with 6 mg/ml samples in 10 mM sodium acetate, pH = 5 in H_2_O:D_2_O 90:10, with 24 scans and 40×128x2048 points in F1: ^1^H: F2: ^15^N and F3: ^1^H dimensions and with a NUS factor of 50%. The spectral width was 12.8 ppm in both proton dimensions and 35 ppm in ^15^N dimension. The inter-scan delay (*d_1_*) was 1 s. The spectra were measured at 25 and 45 °C and at each temperature a mixing time of 80 and 120 ms was used. The measurement time of each spectrum was 21.9 hours. The average volume of the 3% of the intra-residue stronger peaks was used to set the reference distance of 2.5 Å. Other proton-proton distances were calculated using the I∼r ^−6^ relation (31).

### NMR relaxation

^15^N relaxation experiments based on the ^15^N-HSQC detection (30, 76) were carried out at 25 °C and 45 °C at a 6 mg/ml protein concentration. Each spectrum was measured with 8 scans and 128×2048 points in the ^15^N and ^1^H dimensions. The Longitudinal relaxation rates (R_1_) were measured (sequence *hsqct1etf3gptcwg3d*) exploring 12 points of the recovery delay between 2 ms and 1.2 s. Transverse relaxation rates (R_2_) were measured (sequence *hsqct2etf3gptcwg3d*) exploring 12 points of the CPMG duration between 15.6 and 343 ms.

For determining ^15^N_*R_1_* and ^15^N_*R_2_*, the intensity of a given cross peak in the N-HSQC correlation at each relaxation delay *τ*, *I(τ),* was measured and the curve was non-linearly fit to the mono-exponential equation Eq. 3 to determine *I_0_* and *R_x_*.

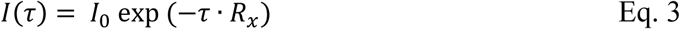

Where *R_x_* refers to either ^15^N_*R_1_* or ^15^N_*R_2_*, and *I_0_* is the initial intensity at *τ* =0.

Steady state ^15^N{^1^H} heteronuclear NOE (^15^N_NOE) (sequence *hsqcnoef3gpsi*) was measured by acquiring two interleaved spectra collected with (*S^sat^*) and without (*S^0^*) an initial proton saturation period (3 s) during the 5 s recycle delay. The ^15^N_NOEs were determined by measuring the intensity of a given cross peak in the saturated (*S^sat^*) and reference (*S^0^*) spectra using Eq. 4.

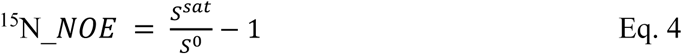

### NMR diffusion experiments (DOSY)

Diffusion-Ordered SpectroscopY (DOSY) spectra were measured with the Oneshot experiment (77). The sequence was adapted to incorporate a watergate 3-9-19 scheme for the strong suppression of the water peak. The diffusion delay Δ was 100 ms; the bipolar gradient pulses (δ) used to encode/decode diffusion have a total duration of 4 ms with a smoothed squared shape. Their power was varied linearly between 2.5 and 50.3 G/cm to detect 30 points in the diffusion dimension with 24 scans per point. The inter-scan delay (d_1_) was 1 s and the measurement time is 27 min. Each DOSY spectrum was processed witn MestreNova and the most-intense proton signals of the protein in the region 0 to 4.5 ppm were integrated and non-linearly fitted to the single-exponential Stejskal-Tanner equation governing the experiment to determine the self-diffusion coefficient (D) (Eq. 5).

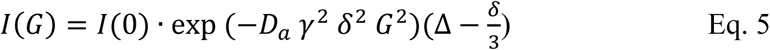

Where I(0) is the original intensity at zero gradient, D is the self-Diffusion coefficient (m^2^·s^−1^), G = gradient strength varied during the experiment (G/m), γ= magnetogyric ratio of the proton (26752.2205 G-1 s-1), δ is the gradient pulse duration and Δ is the diffusion delay. The molecular weight (M) (in g/mol) was estimated from the D value (in m^2^·s^−1^) using the empirical relationship for globular proteins valid from 4 to 120 kDa (78), (Eq. 6).

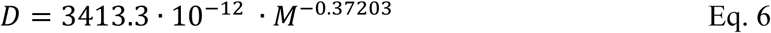

Upon estimating, from Size Exclusion Chromatography (SEC) that in the post-70 sample there was a mixture of two discrete monomeric and oligomeric species (see Results section), we fitted the DOSY intensity data to a double-exponential model (42) (Eq. 7):

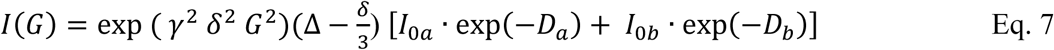

where Dₐ and D_b_ are the diffusion coefficients of the monomer and oligomer, respectively, I₀_a_ and I₀_b_ are their initial intensities and weighting factors, and the other terms have the same meaning as in Eq. 5. Dₐ was fixed at 0.96 ± 0.04 × 10⁻¹⁰ m²/s and weighting factors I₀_b_ and I₀_a_ to the SEC-derived 2:3 ratio.

### Far-UV Circular Dichroism (CD)

For thermal unfolding experiments, CD spectra were acquired in a Jasco 1100 spectrometer with temperature control at a 0.7 mg/ml protein concentration, in 10 mM sodium acetate, pH = 5 placed in a 1 mm light-path quartz cuvette. A temperature ramp from 25 to 85 °C was applied, with spectra between 180 and 260 nm acquired every 1 °C. Five spectra were acquired and accumulated for each data point. For comparative spectra at 25 °C, samples were diluted to a protein concentration of 0.1 mg/ml and CD spectra acquired using a Jasco715 spectrometer and 1 mm light-path quartz cuvettes. Twelve spectra were acquired and accumulated.

CD data as a function of temperature was fitted by SciDavis software to a single-state sigmoidal Boltzmann model to calculate the percentage of unfolding (Eq. 8).

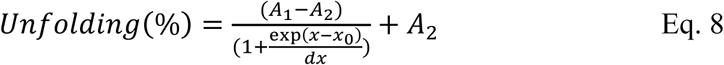

where *A_1_* and *A_2_* are the minimum and maximum value, *x_0_* is the midpoint of the transition and *dx* is the time constant.

### SEC

Samples subjected to different temperatures (*vide supra*) and cooled back to 25 °C. After that they were subjected to SEC using a 90 cm TSK4000SW (7 mm×600 mm) column eluted with 10 mM sodium acetate at 1 ml/min. A_280_ of the effluent was monitored. This column was previously calibrated for molecular mass using low and high molecular mass calibration kits (Amersham). Before each run the column was equilibrated with at least four column volumes of elution buffer. Each run was performed at 20 °C (65).

### Transmission electron microscopy

BVPrP90-231 (10 μl) samples were adsorbed on freshly glow discharged 400 mesh carbon-coated copper grids (Electron Microscopy Sciences), washed with water twice, negatively stained with freshly filtered 2% uranyl acetate for 20 s, air-dried and imaged using a JEOL JEM 1011 electron microscope at 100 KV and a MegaView G3 camera.

### Molecular Dynamics (MD) simulations

MD simulations were performed on the supercomputer Finis Terrae II (CESGA, Santiago de Compostela, Spain). A model representing a BVPrP(90–231) monomer with its 90-120 flexible tail attached to a PrP^Sc^ stack was constructed. We used the experimentally determined structure of BVPrP (121–231) (PDB: 2K56). Since no BVPrP^Sc^ structure has been resolved yet, we used RML MoPrP^Sc^ (PDB: 7td6, ref. 5), as an approximation. To a tetramer of this PrP^Sc^, we added a 93-120 MoPrP rung to which we manually attached the 121-231 FD of BVPrP(121–231). It should be noted that the 93-120 sequences of MoPrP and BVPrP differ in residue 109 only: L in mouse and I/M in BV (I in the construct used in this study). Motions of the PrP^Sc^ tetramer were restricted, to simulate the situation of a hypothetically much more stable >>4-mer stack, and also because at this point only the trajectory of the attached PrP^C^ FD interests us. However, a recent MD study has shown that the surface of a PrP^Sc^ stack does have fluctuations that should be taken in consideration in future studies (79). The resulting structure underwent energy minimization and equilibration using a standard protocol in GROMACS-2022, employing the V-rescale thermostat to maintain a constant temperature and the Parrinello-Rahman algorithm for pressure coupling (80). Molecular dynamics (MD) production simulations were carried out in the NPT ensemble, maintaining the same conditions used during equilibration. One run of 200 ns was performed at 310 K and three independent runs of 300 ns each were done at 400 K.

In parallel we carried out MD simulations of BVPrP^C^(90–231) with its tail free, just as in our experiments. In this case the model was constructed by adding to the BV121-231 structure (PDB: 2K56) the flexible N-terminal region. This N-terminal tail was modeled based on the solution NMR structure of the human PrP molecule (PDB: 7FHQ), which includes residues 91-120. Moreover, residue 90 was added to complete the sequence. To match the BV sequence, residues N97, I109, and V112 were mutated accordingly. The topology was generated using the CHARMM36 force field, which is specifically optimized to accurately model both structured and unstructured regions of proteins. The structure underwent energy minimization and equilibration (*vide supra*). MD production simulations were carried out as detailed for the model described above. Three independent 200 ns production runs were performed at 400 K, and two at 310.

## Acknowledgments

We thank Simone Hornemann, from University Hospital, Zürich, for sharing chemical shift data, and Albina Román-Castro and María José Pazos-Guldrís, from the University of Santiago de Compostela, for TEM imaging.

## Funded by

Spanish National Research Agency (AEI), Spanish Government, co-financed by the European Regional Development Fund-ERDF (grant numbers: PID2020-117465GB-I00 to JRR and PID2024-160022OB-I00 to JC), JPND-Instituto Carlos III (Grant number AC23_2/00049) and Xunta de Galicia ED431C 2024/11. RVC is a pre-doctoral fellow of the Xunta de Galicia. EB and MR were supported by grants from Fondazione Telethon (GGP20043, GMR24T2072). CIQUS is a “Singular Center” (grant ED431G 2023/03, partially funded by the EU ERDF).

## Supplementary Material

**Suppl. Fig. S1.**
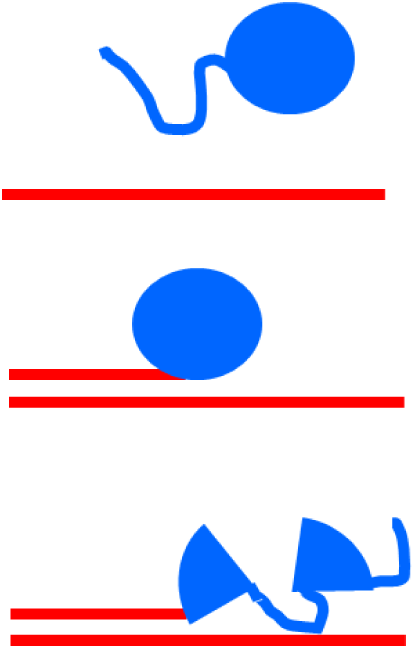
The PrP^C^ folded domain must unfold before it can be templated to refold to the PrP^Sc^ conformation. Top: PrP^C^ (blue) meets PrP^Sc^ (red); its disordered ∼90-120 segment is immediately trapped and converted (middle). However, the PrP^Sc^ surface is a relatively passive template and in order to be refolded the FD of PrP^C^ must first unfold (bottom).

**Suppl. Fig. S2.**
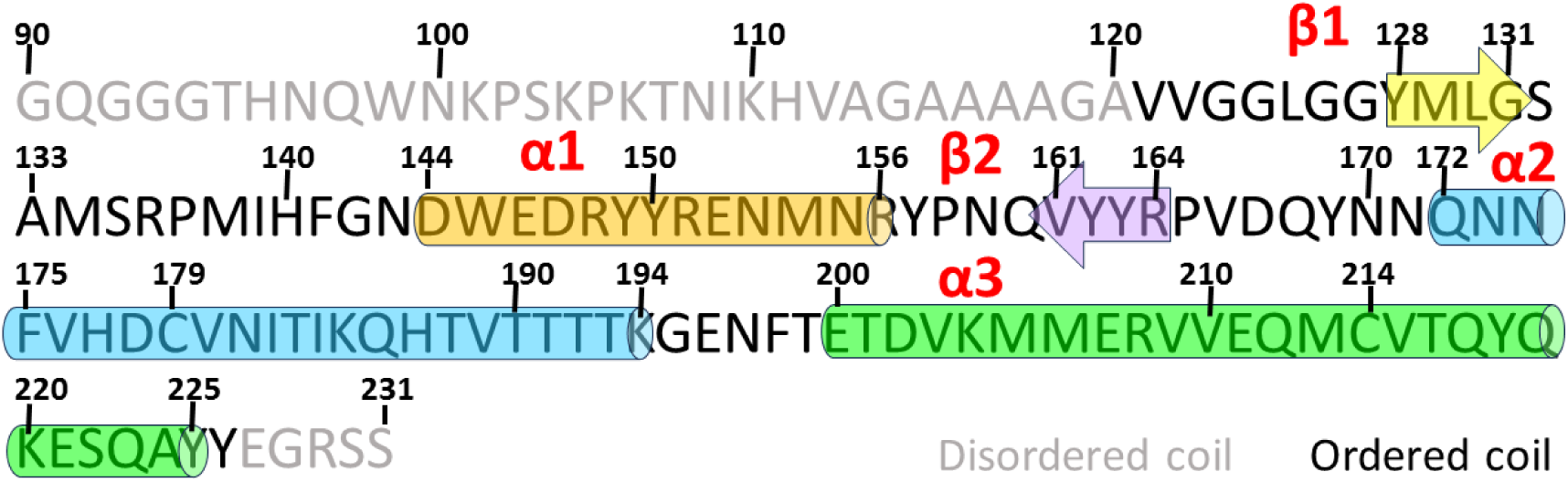
Cartoon showing the sequence of BVPrP90-231 with its secondary structure elements. Based on ref. 12.

**Suppl. Fig. S3.**
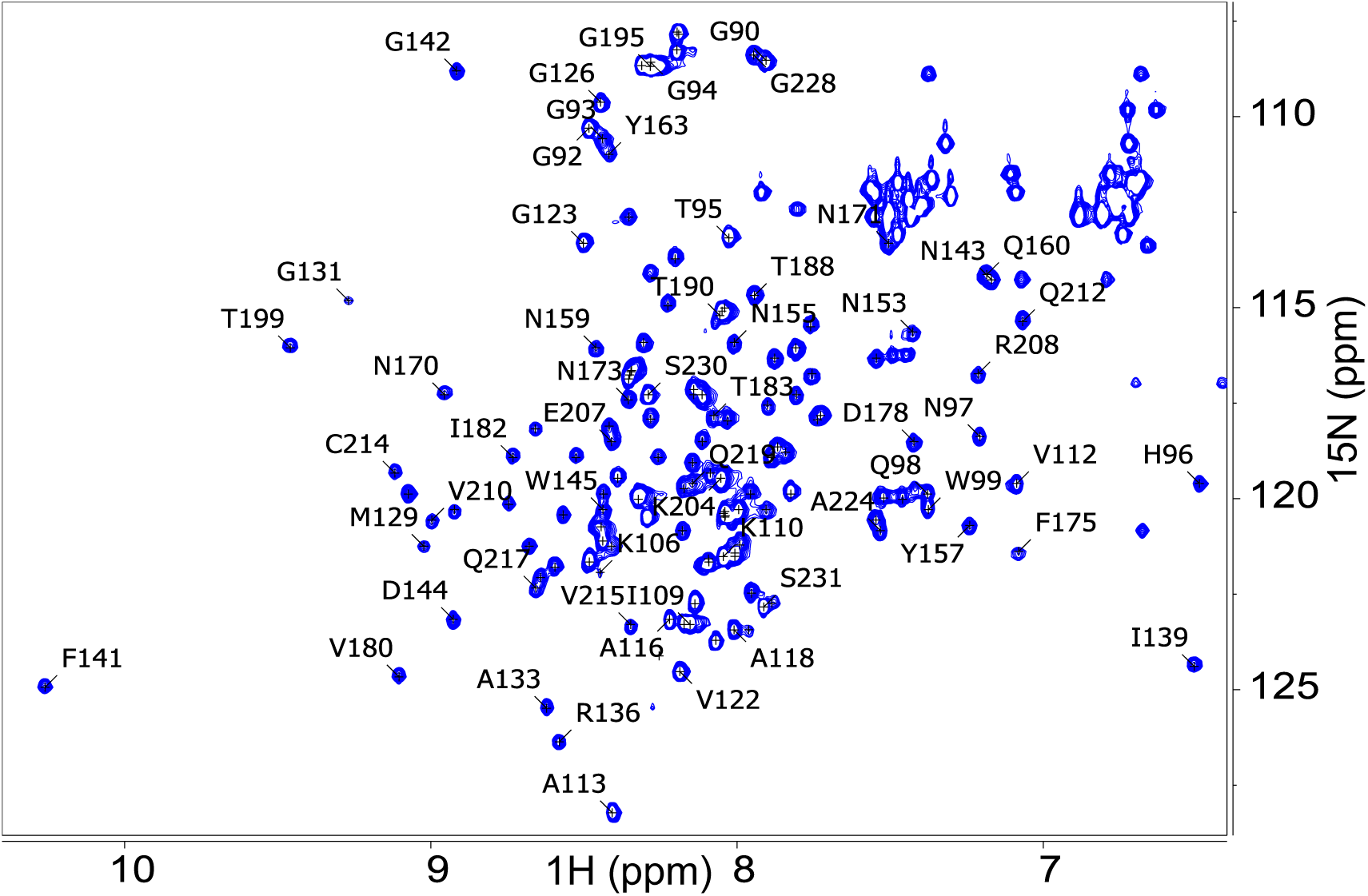
^1^H-^15^N HSQC spectrum of BVPrP^C^(90–231) with assignments (25 ° C, 750 MHz). See main text for details. The protein dissolved in 10 mM sodium acetate buffer, pH 5, was supplemented with D2O (10%). The final protein concentration was 6 mg/ml.

**Suppl. Fig. S4.**
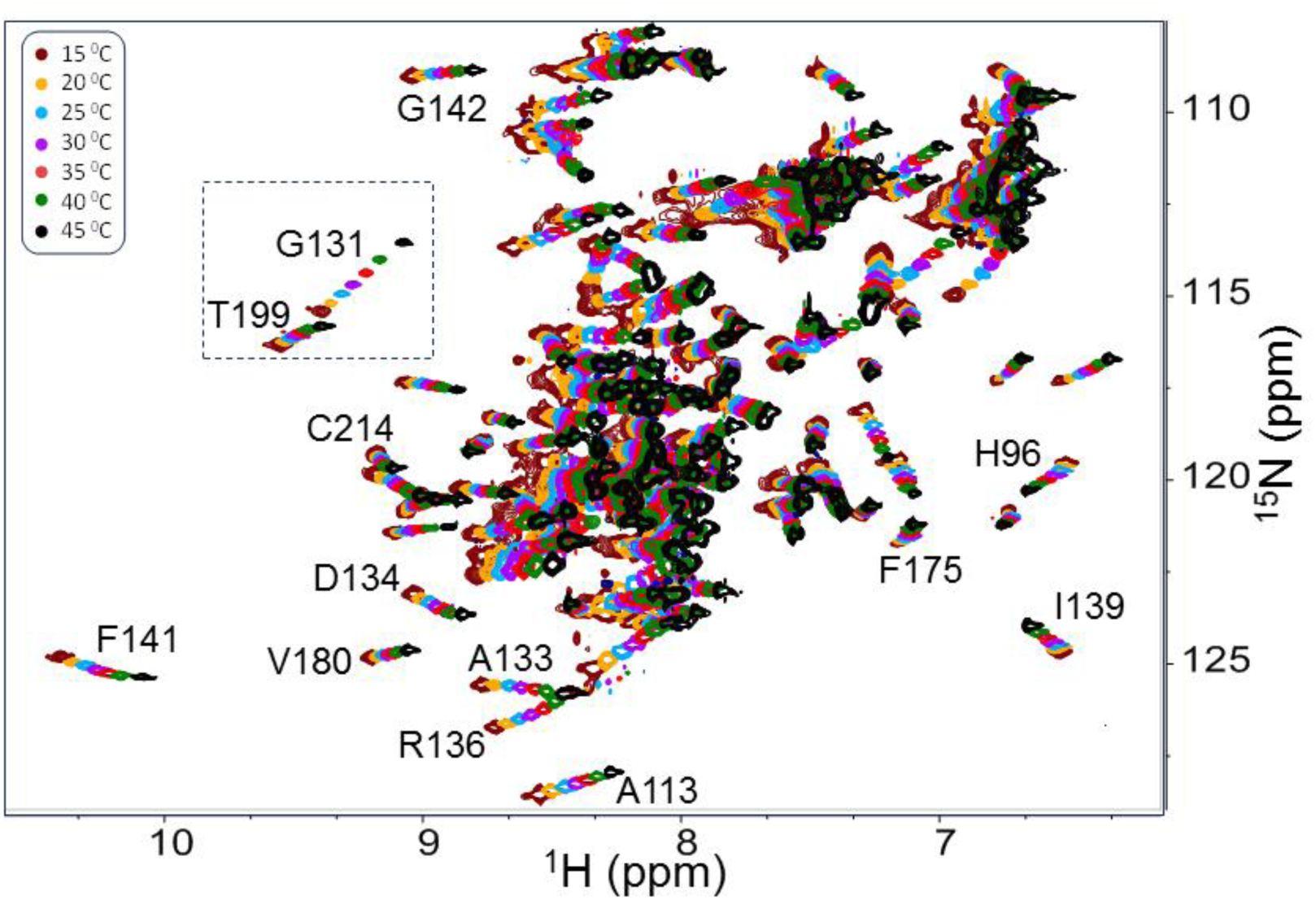
Temperature-induced chemical shift drift of BVPrP^C^(90–231) HSQC signals. The assignment of some of the peaks is shown. The inset highlights a signal (T199) with a linear drift with respect to temperature and another signal/one (G131) with a non-linear drift.

**Suppl. Fig. S5.**
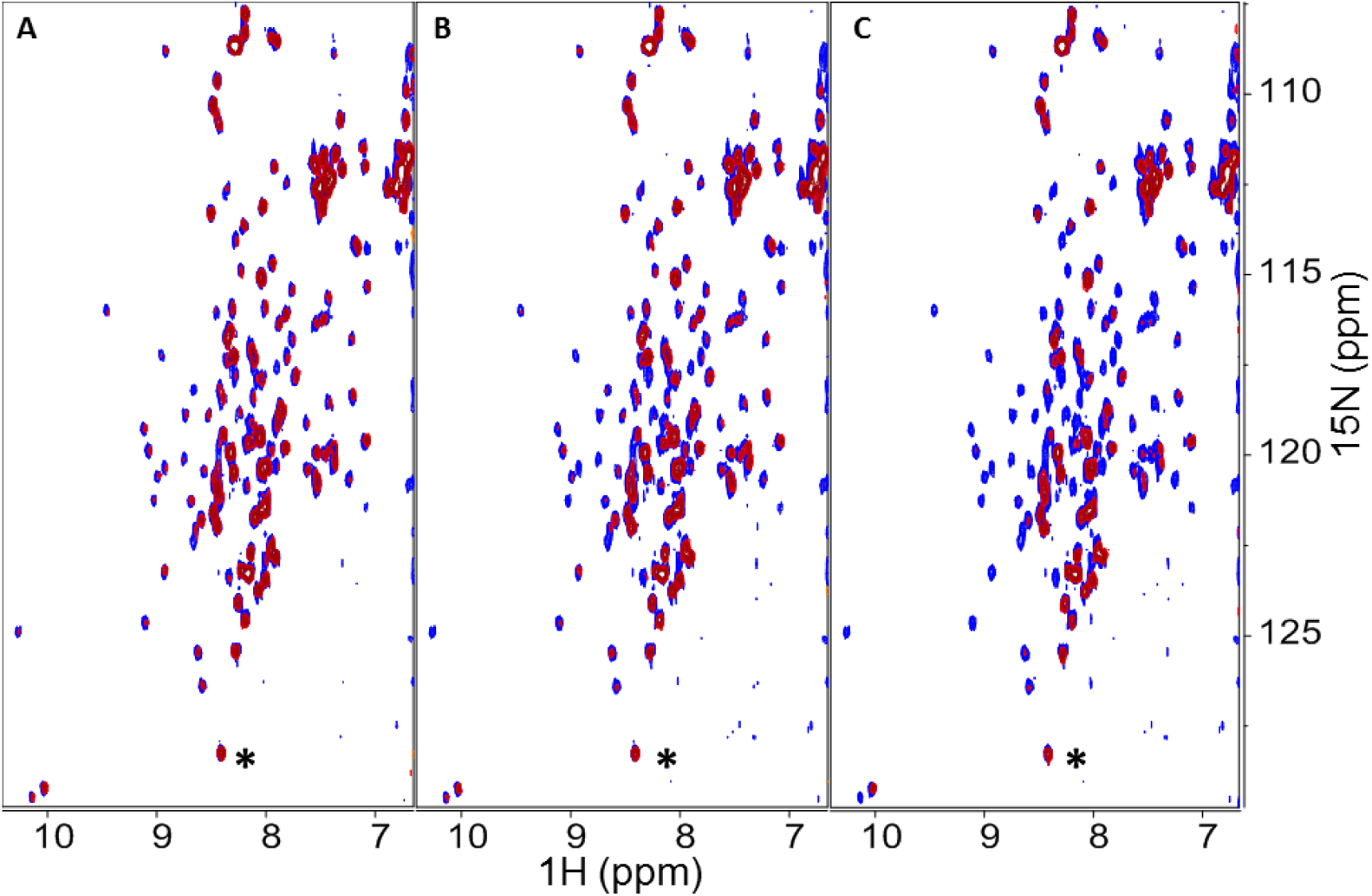
Reversibility/irreversibility of changes in ^15^N-HSQC spectrum of BVPrP^C^(90–231) upon heating. Superimposition of spectra recorded at 25 °C, of the fresh unheated sample (blue) and of the same sample after heating at the indicated temperatures for 1 hour and then cooling at 25 °C (red). *(A)* Heating at 50 °C. Heating at 55 °C *(C)* Heating at 60 °C. The spectra are represented at ca. the same noise level respect to the signal labelled with an asterisk.

**Suppl. Fig. S6.**
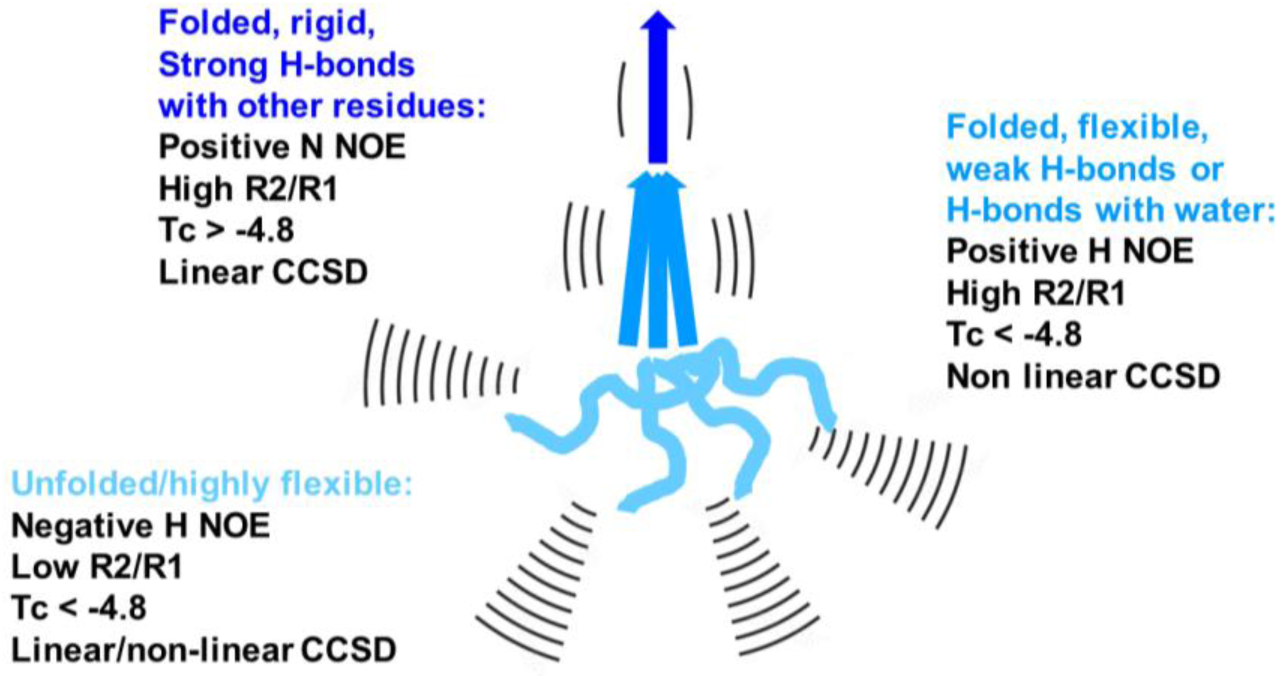
Cartoon illustrating the observables depending on the temperature dependence of amide H-N chemical shifts analyzed though different NMR experiments and used to evaluate flexibility of different regions of a protein. In the present study they were used to identify regions of BVPrP^C^(90–231) likely involved in early unfolding events.

**Suppl. Fig. S7.**
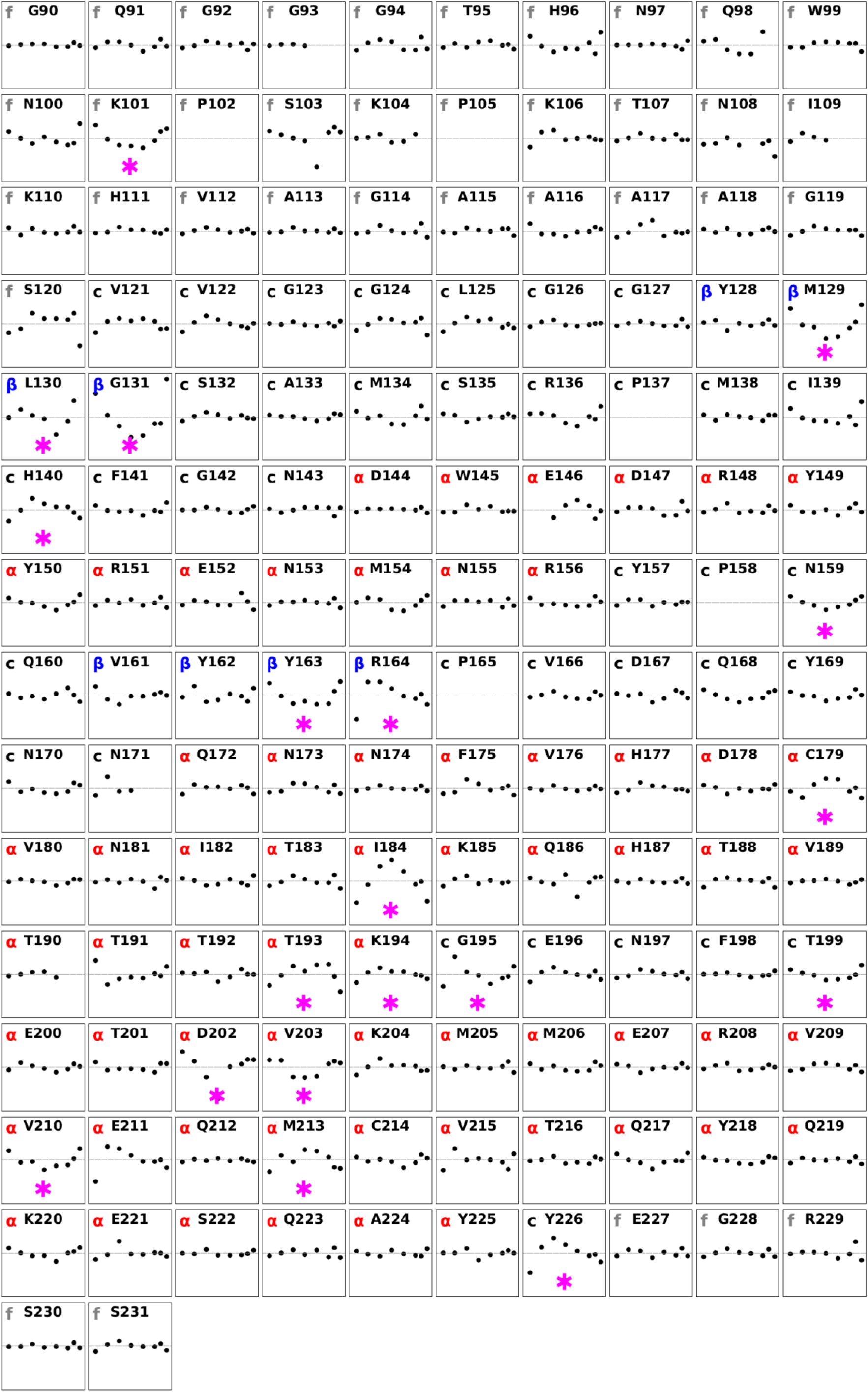
Plot of CCSD residual error for backbone amides of individual BVPrP^C^(90–231) amino acid residues *vs*. temperature. The CCSD values for each residue are calculated by Eq. 1 through peak tracking across the series of VT N-HSQC spectra. They were fitted to a linear regression model with Eq. 2. Each plot represents the residual error (defined as CCSD - CCSD^calc^) against temperature (38, 39). In each plot the amino acid is indicated, and the letter code (in gray) refers to f=flexible (disordered), α=helix, β=sheet or c=coil. The magenta asterisk denotes a residue with non-linear variation of their chemical shifts with temperature.

**Suppl. Fig. S8.**
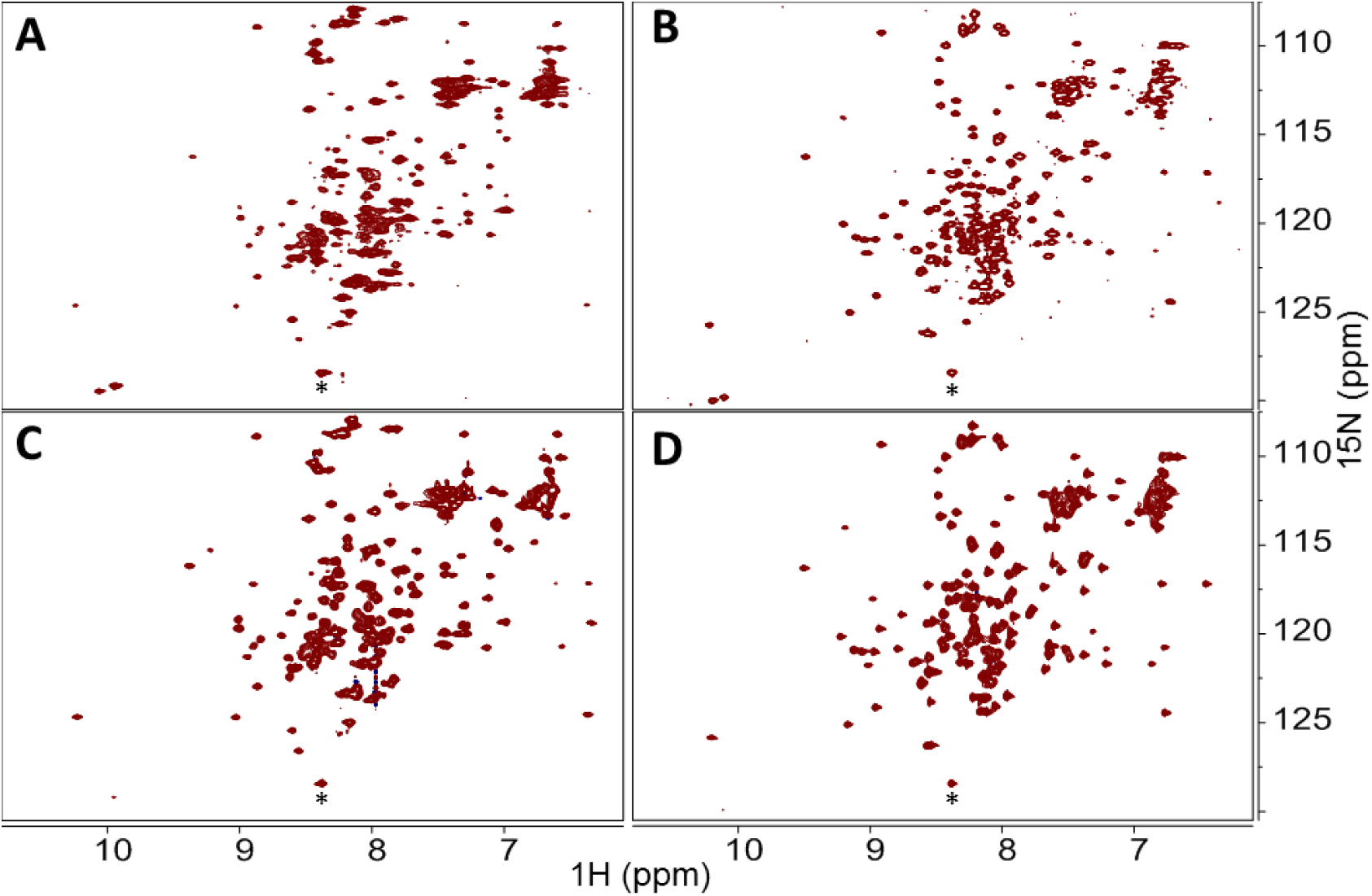
A more concentrate sample (6 mg/mL) behaves like the 1.4 mg/mL sample in the 15-45 °C temperature range. N-HSQC spectra of a BVPrP^C^(90–231): *(A)* 1.4 mg/mL sample at 15 °C. *(B)* 1.4 mg/mL sample at 45 °C. *(C)* 6.0 mg/mL sample at 15 °C. *(D)* 6.0 mg/mL sample at 45 °C. The spectra are represented at ca. the same noise level and the chemical shifts were referenced to the signal marked with the asterisk.

**Suppl. Fig. S9.**
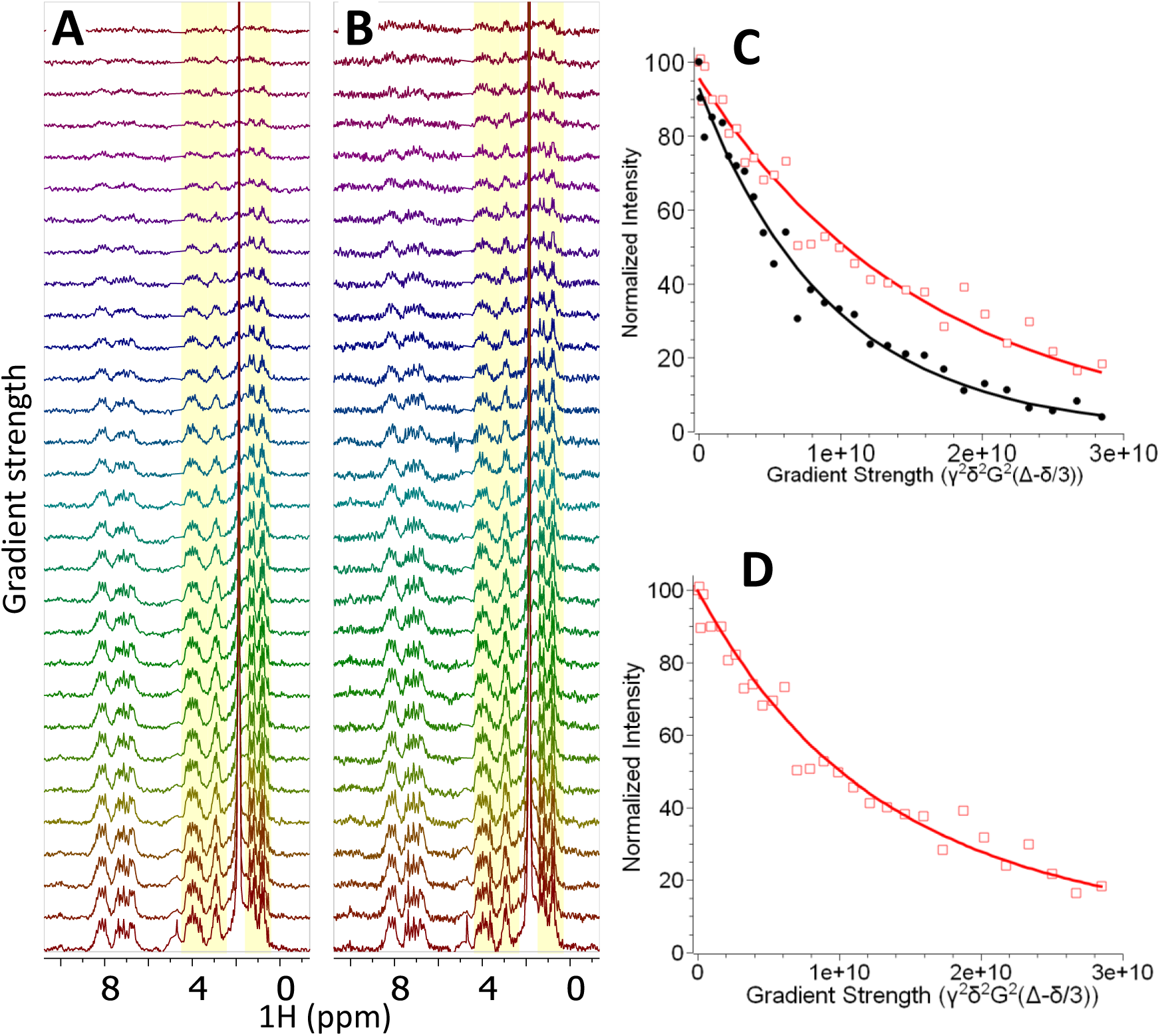
DOSY analysis supports that BVPrP^C^(90–231) undergoes irreversible oligomerization at temperatures above 50 °C. *(A)* DOSY spectrum of a fresh sample at 25 °C (1.4 mg/ml, pH 5). *(B)* DOSY spectrum at 25 °C of the *post-70 °C* sample. DOSY curves were calculated from the integrals of the regions corresponding to non-water exchangeable protons of the protein, that are highlighted in yellow in panels *(A)* and *(B). (C)* Plot of DOSY integral versus gradient strength. Black circles represent the fresh sample, and open red squares represent the post-70 °C sample. Solid lines show fits to the mono-exponential Stejskal-Tanner equation (Eq. 5), yielding diffusion coefficients (D) of 0.96±0.04 × 10⁻¹⁰ m²/s for the fresh sample and 0.63±0.04 × 10⁻¹⁰ m²/s for the post-70 °C sample. The latter D value likely reflects a weighted average of monomeric and oligomeric species that are present in the post-70 °C sample. *(D)* Plot of DOSY integral versus the gradient strength of the post-70 °C sample. The data was fitted to a double-exponential equation analogue of Eq. 5 assuming 0.96 × 10⁻¹⁰ m²/s for D of the monomer species and a ratio of monomer:oligomer 2:3, as deduced from SEC data it provided a D of 0.32±0.03 × 10⁻¹⁰ m²/s for the oligomeric species, which correspond, to a Mw of 270 kDa or ∼16 mers (see Materials and Methods).

**Suppl. Fig. S10.**
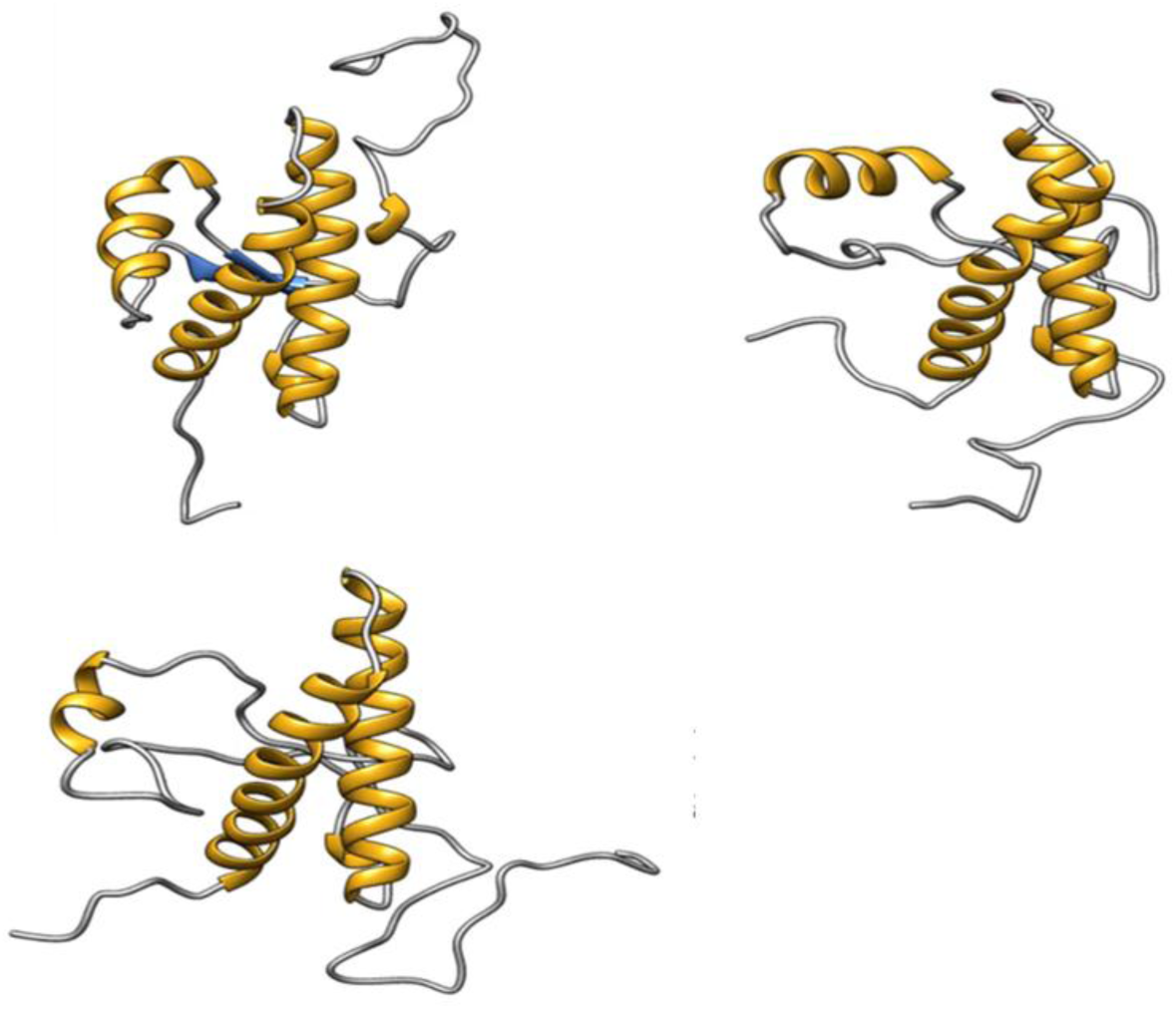
One of two Molecular Dynamics simulation trajectories of evolution of BVPrP^C^(90–231) at 400 K shows separation of α1 from the rest of the FD ensemble. *(A)* t = 0. *(B)* t = 120 ns. *(C)* t = 175 ns. Frames taken from Suppl. Video 6. Note the separation of α1 (t =120 s *vs*. t= 0)

**Suppl. Fig. S11.**
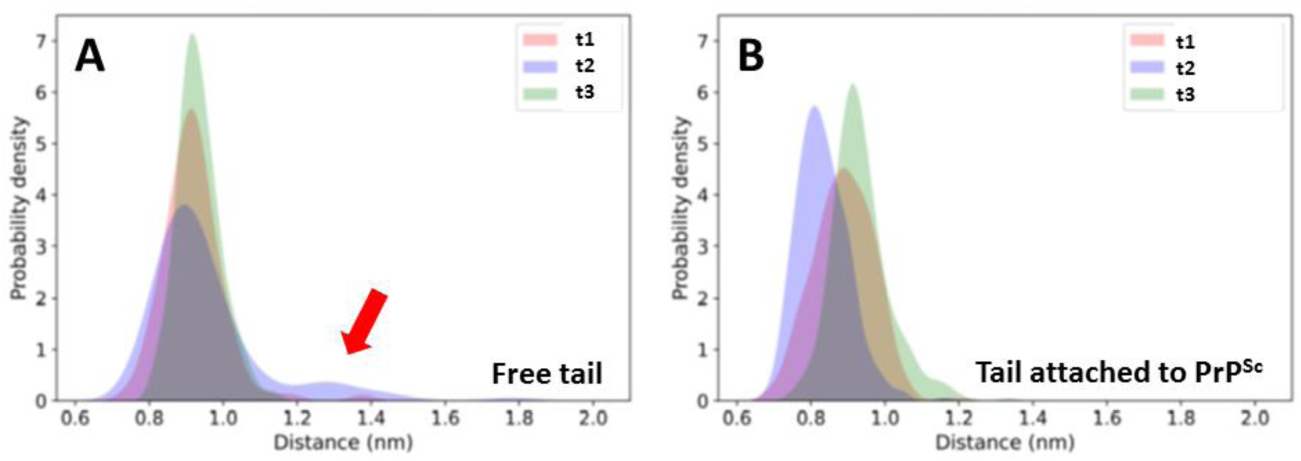
Separation of α1 from the β2-α2-α3 ensemble in some MD trajectories. The plots show probability densities of the distance between α1 and α3, that have an extensive contact surface in BVPrP^C^(90–231). *(A)* BVPrP^C^(90–231) alone. *(B)* BVPrP^C^(90–231) with its 90-120 tail attached to a n immobilized PrP^Sc^ trimer. Three 300 ns trajectories (t1-t3) at 400 K are shown for each model. The red arrow signals a pose with α1 from the β2-α2-α3 ensemble seen in one trajectory.

**Suppl. Fig. S12.**
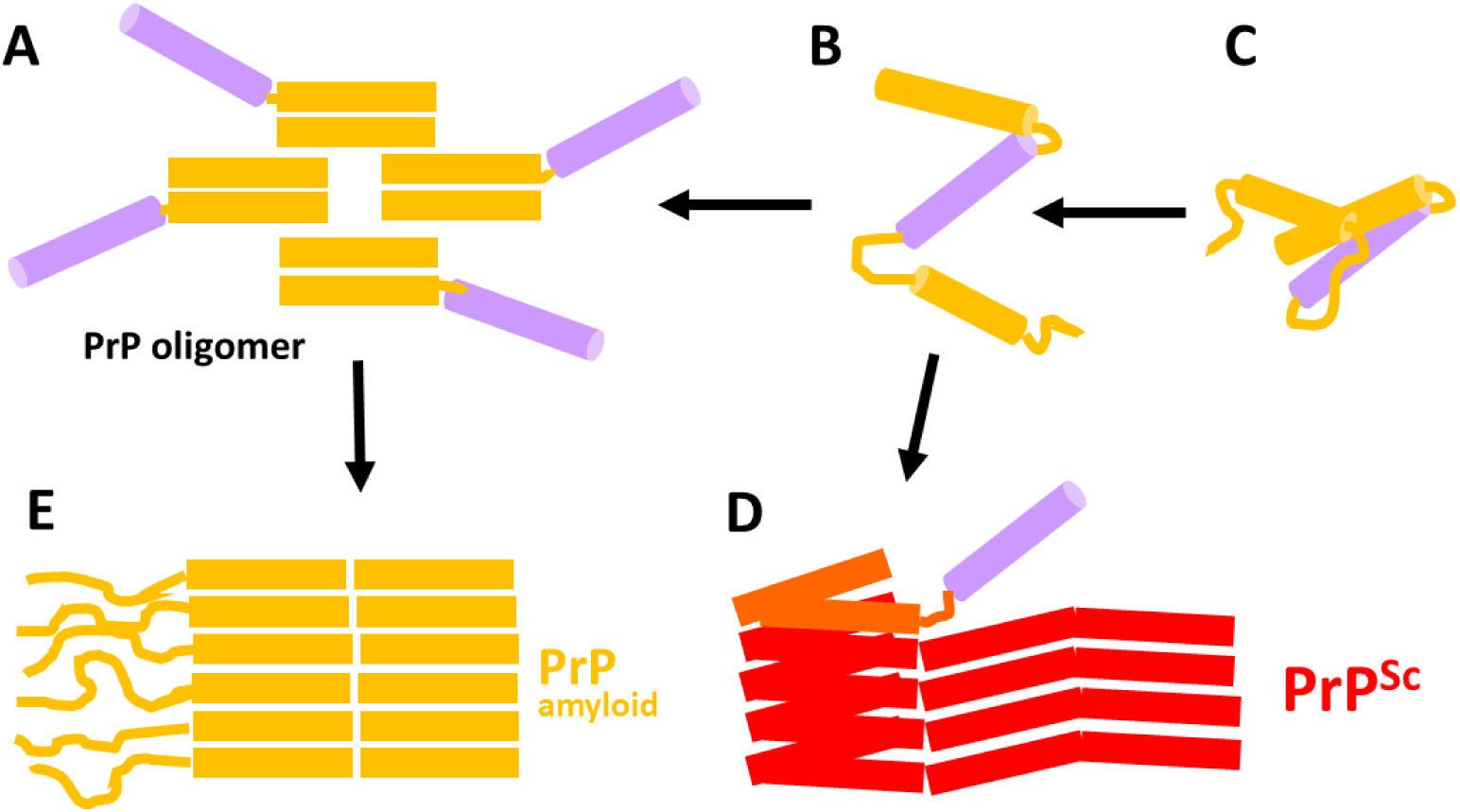
Off-track oligomers provide useful information about an intermediate stage of the unfolding process of BVPrP^C^. The existence of thermically-induced soluble BVPrP oligomers *(A)* with preserved structural elements necessarily requires the existence of a partially unfolded BVPrP species in which such structural elements are also preserved *(B).* Other elements of partially unfolded BVPrP at such stage would have unfolded into an unstable conformation that would lead to oligomerization, likely with simultaneous conversion to β-structure. The B partially unfolded species must in turn originate from an earlier species in which partial unfolding is reversible *(C)*, as seen by reversible changes detected by NMR and described in the main text. Hypothetically, the putative partially unfolded species existing in stage *(B)* and surmised from *(A)*, if attached to the templating surface of a PrP^Sc^ assembly, would not collapse into oligomers, but rather, would collapse onto the templating surface and further template into the PrP^Sc^ conformation *(D)*. This presupposes that similar intermediate unfolding stages occur in the presence and absence of a PrP^Sc^ template. Finally, as seen in many studies, PrP oligomers *(A)* can evolve to PrP amyloids with structures that are different from PrP^Sc^ and exhibit no or little infectivity *(E)*. In conclusion, oligomers are off-track to PrP^Sc^ but provide useful information on the unfolding process of BVPrP, suggesting which BVPrP^C^ structures are more resilient than others and likely to convert at a more advanced stage of unfolding.

**Suppl. Table I:** Evolution of inter-residual NOE distances of BVPrP^C^ 90-231 involving amide protons with other protons upon heating from 25 to 45 ° C.

| Amide proton | proton | 25°C NOE distance (Å) | 45°C NOE distance (Å) | Secondary structural elements involved |
| --- | --- | --- | --- | --- |
| M129 | Y163 NH | 4.74 | 3.50 | $\beta 1 \rightarrow \beta 2$ |
| G131 | V161 NH | 6.22 | 4.86 | $\beta 1 \rightarrow \beta 2$ |
| A133 | Q160 HB | 4.24 | 4.05 | coil $\beta 1-\alpha 1 \rightarrow$ coil $\alpha 1-\beta 2$ |
| M134 | Q160 HA | 3.23 | 3.17 | coil $\beta 1-\alpha 1 \rightarrow$ coil $\alpha 1-\beta 2$ |
| N143 | E146 HB | 4.33 | 4.44 | coil $\beta 1-\alpha 1 \rightarrow \alpha 1$ |
| W145 | R148 HD | 3.54 | 4.24 | $\alpha 1 \rightarrow \alpha 1$ |
| Y149 | M205 HE | 3.86 | 4.31 | $\alpha 1 \rightarrow \alpha 3$ |
| M154 | Y157 HD | 3.42 | 2.84 | $\alpha 1 \rightarrow$ coil $\alpha 1-\beta 2$ |
| Q160 | M213 HE | 4.32 | 3.00 | coil $\alpha 1-\beta 2 \rightarrow \alpha 3$ |
| Y162 | T183 HG | 4.28 | 3.42 | $\beta 2 \rightarrow \alpha 2$ |
| Y162 | I182 HG | 4.89 | 3.72 | $\beta 2 \rightarrow \alpha 2$ |
| V166 | Y225 HD | 4.73 | 4.25 | coil $\beta 2-\alpha 2 \rightarrow \alpha 3$ |
| Q168 | R229 HD | 3.71 | 3.51 | coil $\beta 2-\alpha 2 \rightarrow$ N-ter coil |
| N173 | V176 HG | 4.04 | 4.02 | $\alpha 2 \rightarrow \alpha 2$ |
| D178 | N181 HA | 4.48 | 3.70 | $\alpha 2 \rightarrow \alpha 2$ |
| V180 | V210 HG | 3.92 | 3.35 | $\alpha 2 \rightarrow \alpha 3$ |
| I184 | V210 HG | 4.02 | 3.92 | $\alpha 2 \rightarrow \alpha 3$ |
| T199 | D202 HB | 4.73 | 4.14 | coil $\alpha 2-\alpha 3 \rightarrow \alpha 3$ |
| E207 | V210 HG | 4.47 | 3.37 | $\alpha 3 \rightarrow \alpha 3$ |
| Q212 | V215 HG | 4.80 | 3.82 | $\alpha 3 \rightarrow \alpha 3$ |

**Suppl. Table II:** Summary of the acquisition parameters for measuring 3D NMR spectra in the 750 MHz spectrometer at 25°C^1^.

| Experiment | Total points<br>TD<sup>acq</sup>(<sup>1</sup>H)/<br>TD(<sup>15</sup>N)/TD(<sup>13</sup>C) | NUS<br>factor<br>(%) | sw<sup>1</sup>H/sw<sup>15</sup>N/sw<sup>13</sup><br>C<br>(ppm) | NS | Exp.<br>Time<br>(hours) | Observa<br>tions |
| --- | --- | --- | --- | --- | --- | --- |
| 2D <sup>15</sup>N-HSQC | 2856 x 160 | 100% | 15.9/35/--- | 16 | 0.8 |  |
| 3D HNCO | 2048/40/128 | 30% | 13.9/28/14 | 16 | 8.2 |  |
| 3D best-HNCO | 698/64/128 | 34% | 13.9/35/14 | 16 | 6.2 |  |
| 3D best-HN(CO)CA | 2082/64/128 | 50% | 13.9/28/30 | 16 | 11.7 |  |
| 3D HNCA | 1306 x 64 x 128 | 40% | 4.35/35/30 | 24 | 41.6 |  |
| 3D best-HNCA | 2854 x 64 x 128 | 36% | 15.9/35/80 | 16 | 16.3 |  |
| 3D best-intra-HNCA | 2082/64/128 | 50% | 13.9/28/30 | 16 | 11.9 |  |
| 3D intra-HNCACB | 2082/48/128 | 25% | 13.9/28/80 | 24 | 12.9 |  |
| 3D best-intra<br>HNCACB | 2082/40/128 | 40% | 13.9/28/80 | 32 | 12.0 |  |
| 3D best-intra<br>HNCACB | 2082/64/128 | 50% | 13.9/28/80 | 24 | 12.2 |  |
| 3D best-HNCACO | 2082/64/128 | 50% | 13.9/28/14 | 16 | 11.7 |  |
| 3D HNHA | 2048 x 128 x 128<br>(<sup>1</sup>H) | 40% | 13.9/35/13.9 | 16 | 34.6 |  |
| 3D<br>HNCAHA | 4096/128/80 | 25% | 10.9/35/30 | 16 | 26.9 |  |
| 3D HCCH-TOCSY | 2048 x 64 (<sup>13</sup>C) x 128 | 50% | 13.9/80/80 | 16 | 21.4 | DIPSI-2<br>mix.<br>12ms |
| 3D <sup>15</sup>N<br>TOCSY-HSQC | 2380 x 100 x 128 | 30% | 15.9/35/14 | 24 | 34.7 | DIPSI-2<br>mix.<br>30ms |
| 3D <sup>15</sup>N<br>TOCSY-HSQC | 2380 x 48 x 128 | 30% | 15.9/28/14 | 24 | 15.0 | DIPSI-2<br>mix.<br>60ms |
| 3D <sup>15</sup>N<br>NOESY-HSQC | 2048 x 64(<sup>15</sup>N) x 450 | 40% | 12.8/28/12.8 | 16 | 63.4 | mix.<br>60ms |
| 3D <sup>15</sup>N<br>NOESY-HSQC | 2048 x 40(<sup>15</sup>N) x 128 | 50% | 12.8/28/12.8 | 24 | 21.9 | mix.<br>80,120m<br>s |
| 3D <sup>13</sup>C<br>NOESY-HSQC | 2048 x 64(<sup>13</sup>C) x 450 | 40% | 12.8/80/12.8 | 16 | 63.0 | mix.<br>60ms |
| 3D <sup>13</sup>C<br>NOESY-HSQC | 2048 x 64(<sup>13</sup>C) x 128 | 40% | 12.8/80/12.8 | 40 | 46.5 | mix.<br>80ms,<br>120ms |

<sup>1</sup>These spectra were used for signal assignment of <sup>1</sup>H, <sup>13</sup>C, and <sup>15</sup>N resonances with the sample of BV PrP<sup>C</sup>(90–231) prepared at 6 mg/mL pH=6.0. The NUS factor represents the percentage of hypercomplex acquisition points finally collected in the indirect dimensions respect to the total.

